# Pathological characteristics of pulmonary toxicity in F344 rats exposed by inhalation to cross-linked water-soluble acrylic acid polymers

**DOI:** 10.1101/2021.11.13.468475

**Authors:** Shotaro Yamano, Tomoki Takeda, Yuko Goto, Shigeyuki Hirai, Yusuke Furukawa, Yoshinori Kikuchi, Kyohei Misumi, Masaaki Suzuki, Kenji Takanobu, Hideki Senoh, Misae Saito, Hitomi Kondo, Yoichiro Kobashi, Kenzo Okamoto, Takumi Kishimoto, Yumi Umeda

**Affiliations:** Japan Bioassay Research Center, Japan Organization of Occupational Health and Safety, Hadano, Kanagawa 257-0015, Japan; Department of Pathology, Tenri Hospital, Tenri, Nara 632-8552, Japan; Department of Pathology, Hokkaido Chuo Rosai Hospital, Japan Organization of Occupational Health and Safety, Iwamizawa, Hokkaido 068-0004, Japan; Director of Research and Training Center for Asbestos-Related Diseases, Okayama, Okayama 702-8055, Japan

**Author notes:** Correspondence **Corresponding Author**, Shotaro Yamano, Japan Bioassay Research Center, Japan Organization of Occupational Health and Safety, Hadano, Kanagawa 257-0015, Japan, Tomoki Takeda, Japan Bioassay Research Center, Japan Organization of Occupational Health and Safety, Hadano, Kanagawa 257-0015, Japan. These authors contributed equally (Shotaro Yamano, Tomoki Takeda).

**Keywords:** cross-linked water-soluble acrylic acid polymer (CWAAP), pulmonary disease, intratracheal instillation, rat, transforming growth factor, whole-body inhalation

## Abstract

**Background:** Recently in Japan, six workers at a chemical plant that manufactures resins developed interstitial lung diseases after being involved in loading and packing cross-linked water-soluble acrylic acid polymers (CWAAPs). Since CWAAPs are not on the list of occupational diseases, the present study examined the lung damage potential of two CWAAPs (CWAAP-A and CWAAP-B) in rats, investigated pathological mechanisms, and established a method to rapidly evaluate the harmfulness of CWAAPs.

**Methods:** Using a whole-body inhalation exposure system, male F344 rats were exposed once to 40 or 100 mg/m^3^ of CWAAP-A for 4 hours or to 15 or 40 mg/m^3^ of CWAAP-A for 4 hours per day once per week for 2 months (a total of 9 exposures). In a separate set of experiments, male F344 rats were administered 1 mg/kg CWAAP-A or CWAAP-B by intratracheal instillation once every two weeks for 2 months (a total of five doses). Lung tissues, mediastinal lymph nodes, and bronchoalveolar lavage fluid were collected and subjected to biological and histopathological analyses.

**Results:** A single 4-hour exposure to CWAAP caused alveolar injury, and repeated exposures resulted in regenerative changes in the alveolar epithelium with activation of TGFβ signaling. During the recovery period after the last exposure, some alveolar lesions were partially healed, but other lesions developed into alveolitis with fibrous thickening of the alveolar septum. Rats administered CWAAP-A by intratracheal instillation developed qualitatively similar pulmonary pathology as rats exposed to CWAAP-A by inhalation. At 2 weeks after intratracheal instillation, rats administered CWAAP-B appeared to have a slightly higher degree of lung lesions compared to rats administered CWAAP-A, however, there was no difference in these lesions of CWAAP-A and CWAAP-B in rats examined 18 weeks after administration of these materials.

**Conclusions:** The present study provides evidence of rat lung pathogenesis after inhalation exposure to CWAAP-A. This study also demonstrates that the lung pathology of rats exposed to CWAAP-A by systemic inhalation was qualitatively similar to that of rats administered CWAAP-A by intratracheal instillation. The use of intratracheal instillation as an adjunct to systemic inhalation is expected to significantly accelerate the risk assessment for a variety of CWAAPs.

## Background

Cross-linked water-soluble acrylic acid polymer (CWAAP) (CAS No.: 9003-01-4) is a type of thickener that improves viscosity and sol-gel stability. Although synthetic superabsorbent polymers (SAP) are also classified as acrylic acid polymers, CWAAP is a different material from SAP in terms of its cross-linked structure and degree of neutralization. CWAAPs have long been used in a variety of products, including cosmetics and pharmaceuticals, because of their low potential for skin and eye irritation. However, in Japan, six workers at a chemical plant that manufactures the resins recently developed interstitial lung diseases after being involved in weighing, packing, and transporting CWAAP [1]. Five of the workers who suffered from lung diseases had a short work history, around two years [1]. Inhalable CWAAPs are not on the list of occupational diseases because pulmonary disorders caused by these materials has not previously been observed. At the request of the Ministry of Health, Labour and Welfare (MHLW), we conducted an accident survey and clinical research of workers involved in the manufacture and transport of CWAAPs, including the aforementioned six patients, in collaboration with the Japan Organization of Occupational Health and Safety. We reported that (1) the clinical and pathological status of the disease in current and past workers could be determined [2], and (2) the individual exposure concentration of respirable particles was as high as 41.8 mg/m^3^ [2]. The MHLW has held five meetings since October, 2018, to discuss respiratory diseases caused by CWAAP dusts: five workers were determined to have work-related illnesses, and these were certified as occupational injuries in April, 2019 [3].

Inhalation exposure to CWAAPs had not yet been reported to cause pulmonary diseases in humans or laboratory animals. As noted above, both CWAAPs and SAPs are hydrophilic acrylic acid polymers, however, their cross-linked structures and degree of neutralization are different, giving them different properties and potentially different effects on living organisms. Nevertheless, it is important to note that based on inhalation test data using laboratory animals Deutsche Forschungsgemeinschaft has set the maximum workplace concentration for SAPs (neutralized cross-linked acrylic acid polymer sodium salt; CAS number 9003-04-7) at 0.05 mg/m^3^ [4]. Importantly, human epidemiological evidence was also assessed, but no effects of SAPs on lung function could be demonstrated.

A recent study suggests that a single intratracheal administration of CWAAP to rats caused transient acute inflammation [5]. However, there is no similar evidence from inhalation exposure of CWAAPs, and in particular, the emergence of chronic pulmonary diseases such as fibrotic changes that occurred in the workers mentioned above has not been reported in animal studies.

The lung disease in workers caused by CWAAPs has not been fully resolved, therefore, to protect the health of workers who work with CWAAPs and to provide appropriate care to patients suffering from respiratory disease potentially caused by CWAAPs, there is an urgent need to elucidate the characteristics and mechanisms of respiratory disease of CWAAPs using animal models. To address this issue, the present study investigated acute and chronic effects of inhalation exposure of rats to CWAAP using a whole body inhalation system.

Interstitial lung disease encompasses numerous lung disorders characterized by inflammation and scarring, i.e., pulmonary fibrosis, that make it hard for the lungs to get enough oxygen, and it is well known that transforming growth factor (TGF) β signaling is important in the progression of pulmonary fibrosis [6]. TGFβ binding to its receptor activates a pathway that results in phosphorylation of the transcription factors Smad2 and Smad3 proteins, which form a heterotrimer with Smad4. This complex migrates to the nucleus where it induces the expression of a number of target genes that promote fibrosis [7]. Therefore, it is very important from a pathological viewpoint to clarify whether CWAAP affects TGFβ signaling and to identify the cells in which TGFβ signaling is activated.

Another important consideration is the large number of CWAAPs with different molecular weights and degrees of cross-linking and the lack of information on their hazard to human health. Thus, establishment of a rapid and simple method to evaluate the risk of different CWAAPs is also very important. This study investigated using intratracheal instillation as an adjunct to inhalation exposure.

## Results

### Repeated systemic inhalation exposure to high concentrations of CWAAP caused pulmonary alveolar damage but not bronchiolar damage

Rats were exposed to CWAAP-A by inhalation at concentrations of 15 and 40 mg/m^3^ for 4 hours per day once per week for 2 months (a total of 9 exposures): 40 mg/m^3^ is the highest exposure concentration found in the workplace [2]. No significant effects on final body weight or general condition were observed (Fig. 1A). Immediately after exposure, there was a statistically significant, concentration-dependent increase in lung and mediastinal lymph node weights compared to the control group (Fig. 1B, C). These increases were still apparent after a 2-week recovery period (Fig. 1B, C). After 18 weeks recovery, the lung and mediastinal lymph node weights were less than the weights after the 2-week recovery period, but the lung weights were still significantly higher in both the 15 and 40 mg/m^3^ groups compared to the control group and mediastinal lymph node weights were significantly higher in the 40 mg/m^3^ group compared to the control group (Fig. 1B, C).

**Figure 1.**
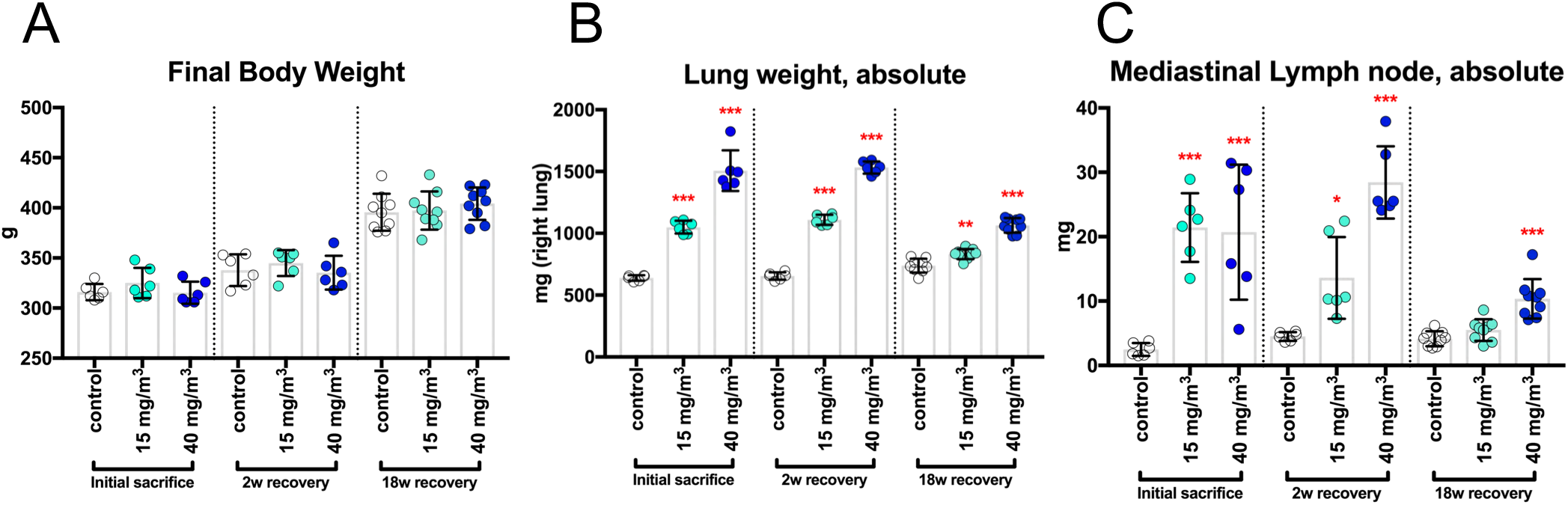
Final body weights and organ weights of the lung and mediastinal lymph nodes of male rats after repeated inhalation exposure to cross-linked water-soluble acrylic acid polymer (CWAAP)-A (15 or 40 mg/m^3^, 4 hours/day, once a week, 9 times). Final body weights (A), right lung weights (B), and mediastinal lymph node weights (C) were measured at each sacrifice. Statistical significance was analyzed using Dunnett’s multiple comparison test compared with age-matched control (sham air) groups: \**p*<0.05, \*\**p*<0.01 and \*\*\**p*<0.001.

Representative images of the lungs are shown in Fig. 2. In the CWAAP-exposed groups, dark red patches, indicative of edema, were observed scattered over the surface of the lung immediately after the end of the exposure period and were still present after the 2-week recovery period (Fig. 2). After the 18-week recovery period, there was recovery of edematous changes, but white spots were observed throughout the lung. In addition to these white spots, distinctive white structures were observed on the cardiac surface on the left lung (Fig. 2, enlarged portion and Table S1).

**Figure 2.**
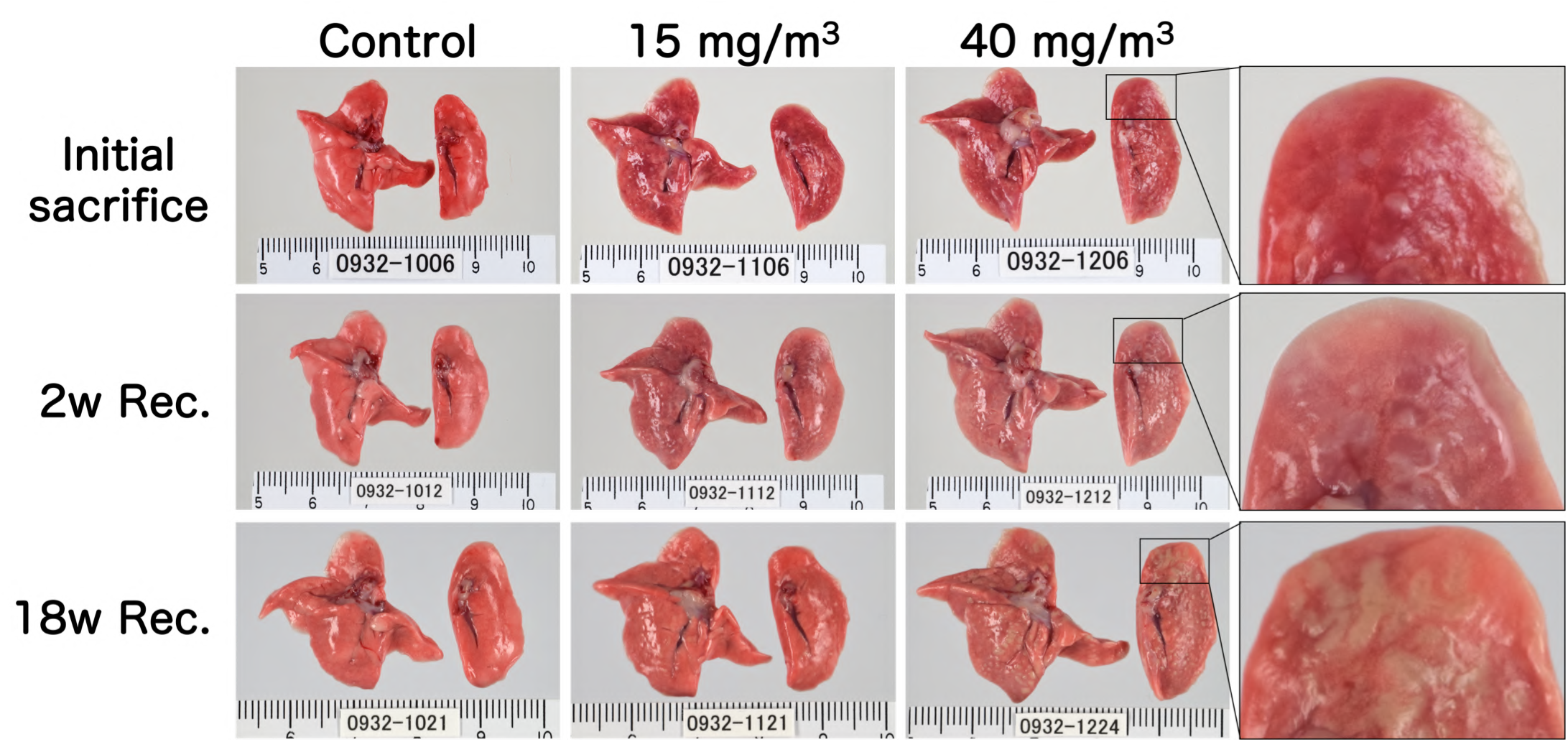
Representative macroscopic photographs of male rat lungs after repeated inhalation exposure to CWAAP-A. High magnification views of the left lung are shown in the right panels. Abbreviations: Rec, recovery; and w: week.

**Table 1.**
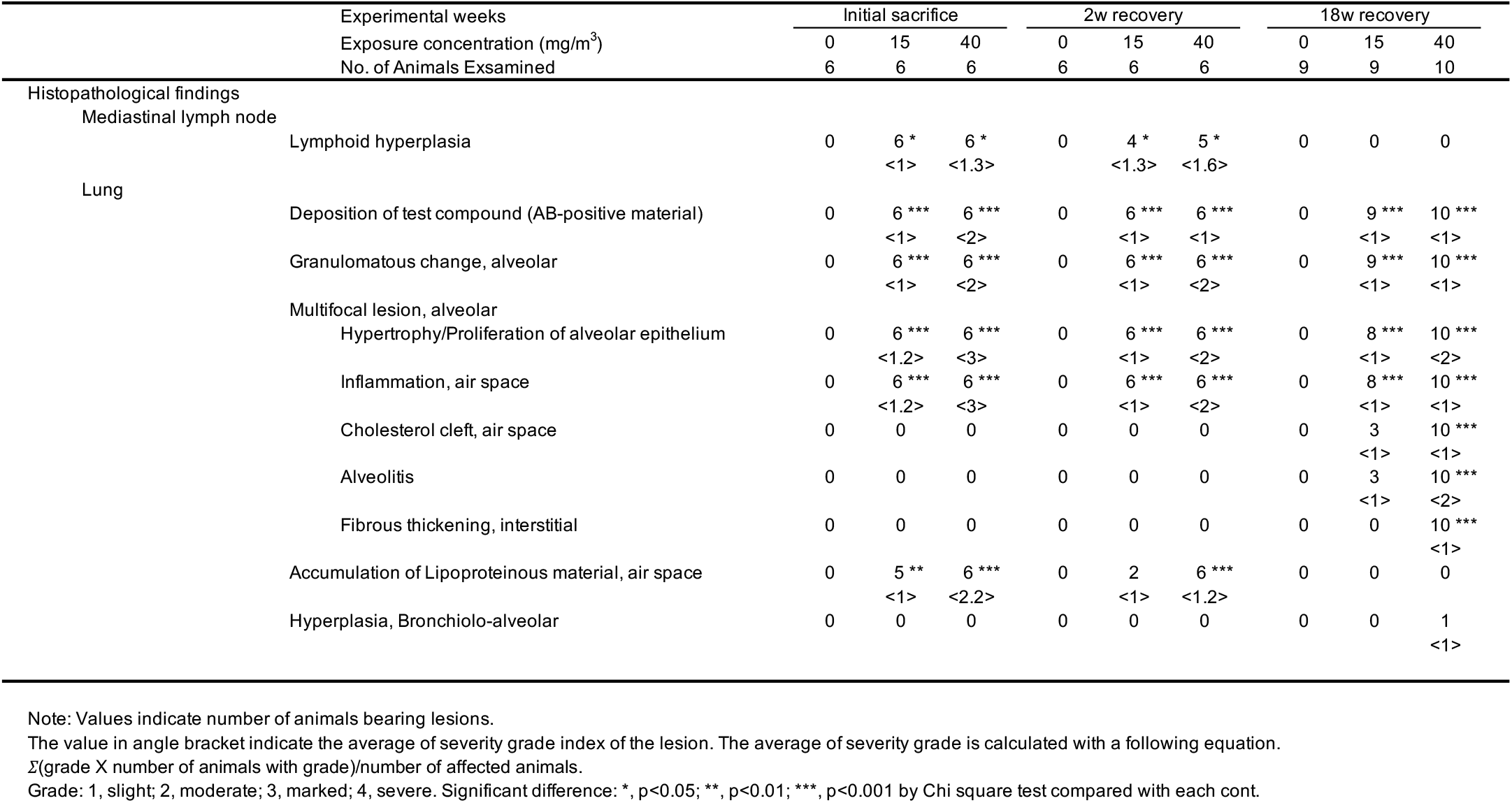
Histopathological findings of the mediastinal lymph node and lung after repeated inhalation exposure to CWAAP-A.

Representative histopathological photographs and cross-sectional images are shown in Fig. S1 (control) and Fig. 3 (CWAAP-A). Fig. 3A shows a representative lung section of a 40 mg/m^3^ exposed rat taken immediately after the end of the exposure period. There were areas of histopathologically observed alveolar proteinosis-like changes that could be identified by Periodic acid Schiff (PAS)-Diastase positive staining: in the area marked hotspot (Fig. 3A upper panel), PAS-Diastase staining was positive (Fig. 3A middle panel), indicating the presence of lipoprotein-like material, which is reminiscent of pulmonary alveolar proteinosis. In addition, multifocal lesions consisting of hypertrophy/proliferation of alveolar epithelium and inflammation were observed as white spots (Fig. 3A lower panel). Hotspot PAS-Diastase positive staining and multifocal lesions (white spots) were also observed in the lungs of rats exposed to 15 mg/m^3^ CWAAP-A.

**Figure 3.**
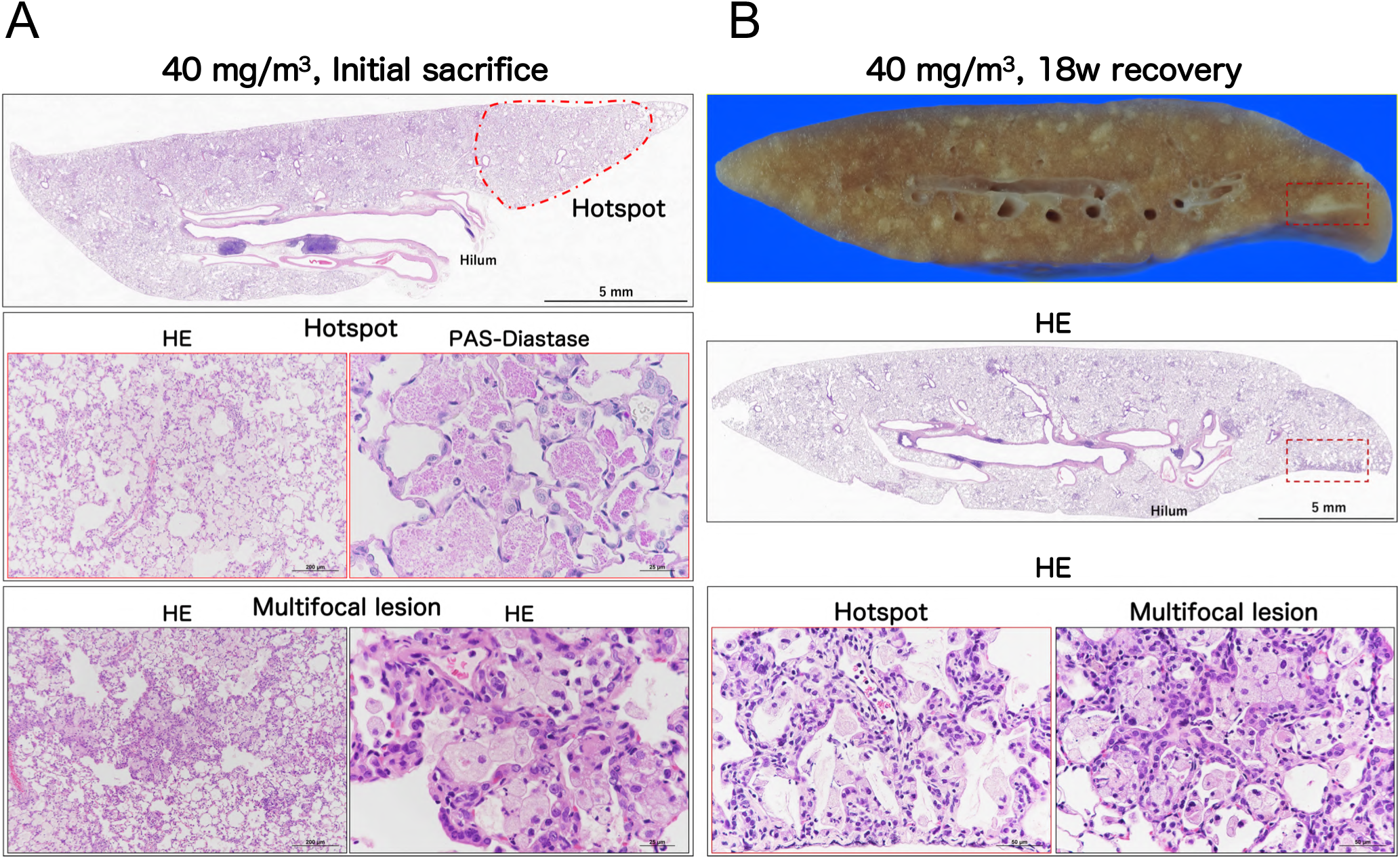
Representative macroscopic and microscopic photographs of rat lungs after repeated inhalation exposure to CWAAP-A. Representative lesions of the rat lung exposed to 40 mg/m^3^ CWAAP-A are shown after hematoxylin and eosin (HE) and Periodic acid Schiff (PAS)-Diastase staining. The boxed areas in the upper and middle panels of 3B outline multifocal lesions after the 18-week recovery period.

Fig. 3 B shows a representative lung section of a 40 mg/m^3^ exposed rat taken after the 18-week recovery period. Prominent white spots (multifocal lesions) were observed in lung cross-sections (Fig. 3B, upper and middle panels). In the hotspot area, a large number of inflammatory cells were observed in the alveolar and perivascular interstitium, spreading to the subpleural area (Fig. 3B, middle and lower left): we diagnosed this as alveolitis. Alveolitis was also observed in multifocal lesions (Fig. 3B, lower right). Alveolitis also developed in the lungs of rats exposed to 15 mg/m^3^ CWAAP-A. These findings indicate that the lesions observed at the end of the exposure period in rats exposed to 15 and 40 mg/m^3^ CWAAP-A continued to develop during the recovery period. In sharp contrast to the alveoli, in the bronchus and bronchiole regions neither epithelial cells nor the surrounding interstitium were prominently affected by CWAAP exposure (Fig. S2).

CWAAP has numerous carboxyl groups (Fig. S3); therefore, to observe the localization of CWAAP in lung tissue we investigated staining methods that in principle react with carboxyl groups. Consequently, we focused on a modified alcian blue staining method, and found a condition in which the mucus of epithelial cells in normal rat lung tissue was not stained, while CWAAP particles were stained blue (Fig. S4). The results of representative alcian blue staining and hematoxylin and eosin (HE) staining in serial sections of a rat lung from the 40 mg/m^3^ CWAAP-A exposed group immediately after the exposure period are shown in Fig. 4. In the lungs of the CWAAP-exposed group, blue-stained areas were observed on the apical cell membranes of the alveolar epithelium, in the alveolar air space, and in the macrophagy cytoplasm. The positive areas were not localized to foci. Within the granulomatous lesions in the alveolar region, prominent blue areas could be observed, but alcian blue staining varied considerably between lesions. These results suggest that alcian blue staining may be useful for the observation of intrapulmonary CWAAP. However, further development of the method is necessary to fully understand the localization of CWAAP.

**Figure 4.**
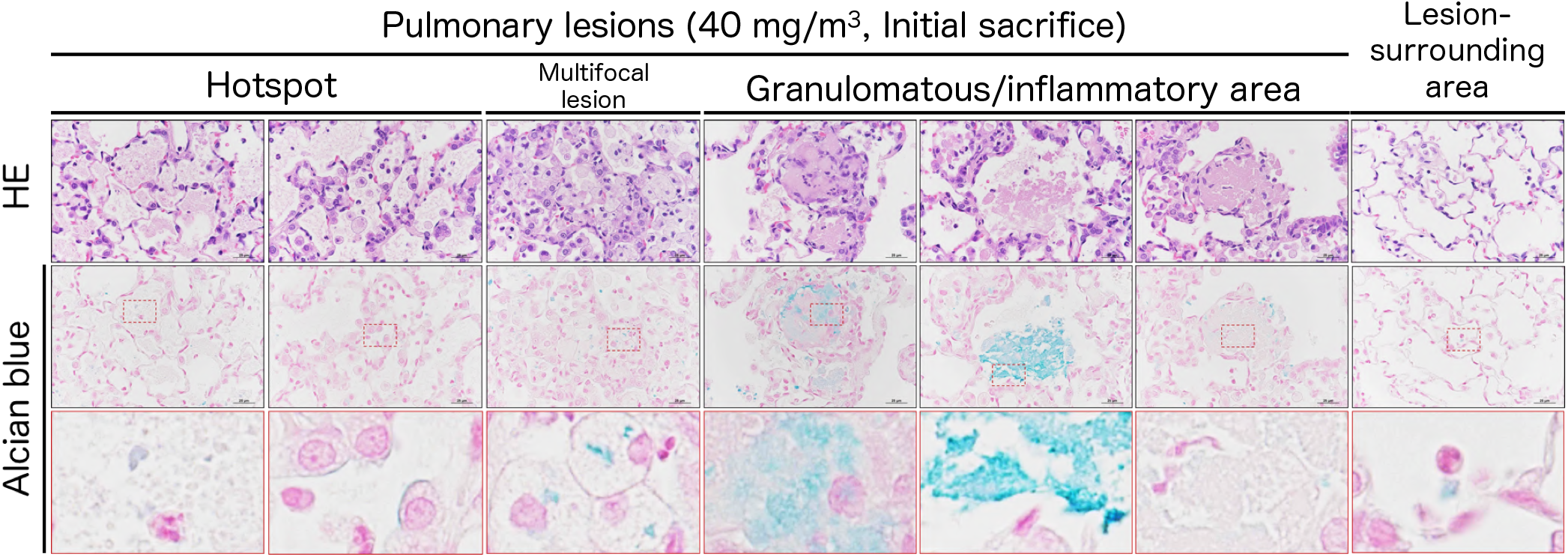
Representative images of alcian blue staining together with HE staining in the lesions and lesion-surrounding areas of the rat lungs after repeated inhalation exposure to 40 mg/m^3^ CWAAP-A.

To examine changes in the fiber volume, Masson’s trichrome staining and the measurement of hydroxyproline levels in the lungs were performed. The results showed that in the lungs of the 40 mg/m^3^ exposure group after the 18-week recovery period there was a marked increase in Masson stain-positive areas in the alveolar septa and perivascular interstitium within the alveolitis (Fig. 5A right panels) compared to the controls and to the rat lung at the end of the exposure period (Fig. 5A left and middle panels). Consistent with this, the hydroxyproline content in the lungs showed significant increases over controls only in the exposed groups sacrificed after the 18-week recovery period (Fig. 5B): there was a concentration-dependent increase in hydroxyproline content in the lungs. These results indicate that prominent fibrous thickening (an increase in collagenous fibers) in the interstitium progressed over time.

**Figure 5.**
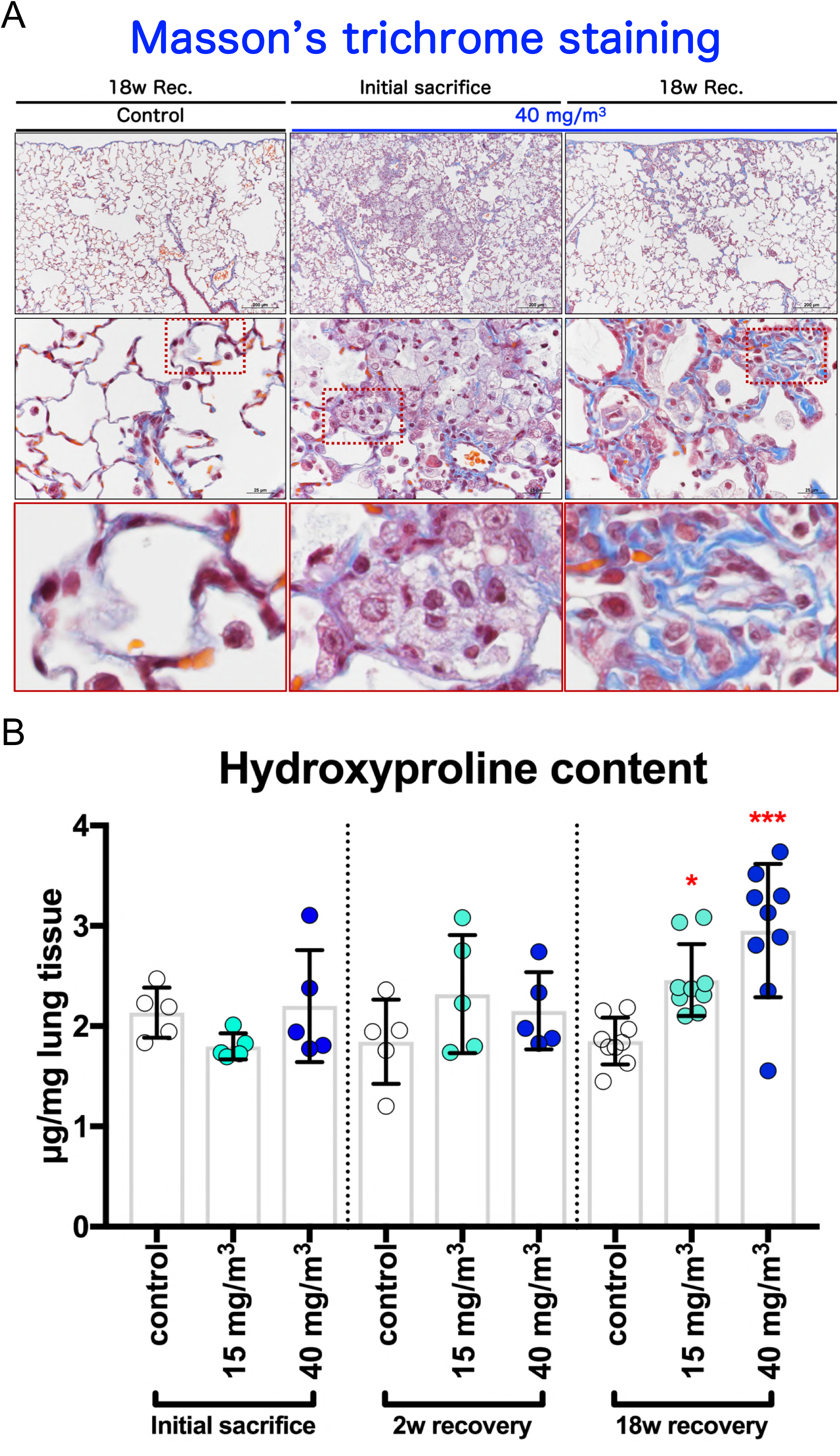
Collagen deposition of the rat lungs after repeated inhalation exposure to CWAAP-A. Representative images of Masson’s trichrome staining (A) and hydroxyproline content (B) in the lung. Dunnett’s multiple comparison test of rats exposed to CWAAP with age-matched control (sham air) groups shows a significant increase in hydroxyproline content after the 18 week recovery period: \**p*<0.05 and \*\*\**p*<0.001.

The histopathological findings in the lung and mediastinal lymph nodes are shown in Table 1. The enlargement of the mediastinal lymph nodes observed in the exposure groups was histopathologically diagnosed as lymphoid hyperplasia. The rats recovered from this condition over time, such that there was no lymphoid hyperplasia observed after the 18-week recovery period. In the lung, granulomatous change, multifocal lesions, and accumulation of lipoproteinous material were observed in the alveolar region in the exposed groups. The multifocal lesions consisted of inflammation in the air space (similar to lipoid pneumonia), and hypertrophy/proliferation of alveolar epithelium in the animals immediately after exposure. In addition, after the 18-week recovery period, histopathological findings included cholesterol cleft in the air space (also known as cholesterol granuloma), alveolitis (also known as interstitial pneumonia), and fibrous thickening in the interstitium.

Interestingly, after the 18-week recovery period, one animal in the 40 mg/m^3^ group showed bronchiolo-alveolar hyperplasia, a pre-neoplastic lesion. Although this lesion was found in only one of the ten animals and is consequently without statistical significance, it was too early to appear as an age-related lesion, suggesting that it may have been caused by CWAAP-A exposure.

Fig. 6 shows the results of the analysis of bronchoalveolar lavage fluid (BALF). In the sham air group, normal macrophages with fine vacuoles were observed. However, in the 40 mg/m^3^ group, a large number of neutrophils, CWAAP deposits, and enlarged macrophages phagocytosing CWAAP were observed both immediately after the exposure period and after the 18-week recovery period (Fig. 6A). The cell numbers found in the BALF are shown in Fig. 6B-D. Lactate dehydrogenase (LDH) activity, a cytotoxicity marker, and surfactant protein-D (SP-D) levels, an interstitial pneumonia marker [8], in the BALF and the plasma are shown in Fig. 6E-G. Immediately after exposure, these all markers were significantly increased compared to the control group in a concentration-dependent manner. This increase was maintained after the 2-week recovery period. However, after the 18-week recovery period, there was a marked decrease in these marker values compared to the values immediately after exposure, and the 15 mg/m^3^ group recovered to the same level as the control group. These results support of the results obtained from histopathological examinations that CWAAP-A caused a multifocal pattern of alveolar lesions, including inflammation in the air space.

**Figure 6.**
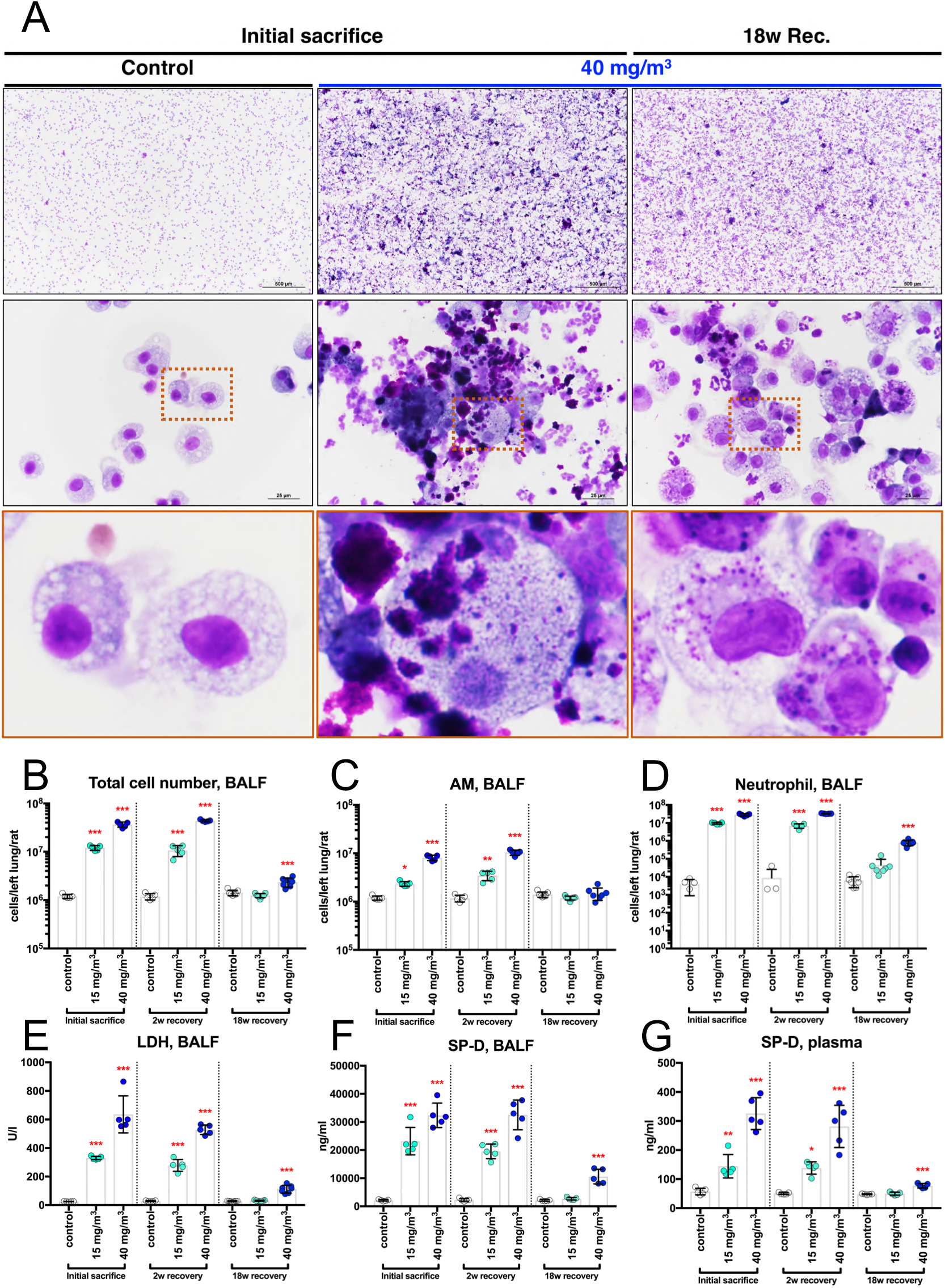
Representative images of the bronchoalveolar lavage fluid (BALF) cytospin cytology (A). Total cell number (B), alveolar macrophage (AM) number (C), neutrophil number (D), LDH activity (E) and surfactant protein-D (SP-D) level (F) in the BALF and SP-D level in the plasma (G). Statistical significance was analyzed using Dunnett’s multiple comparison test: \**p*<0.05, \*\**p*<0.01, and \*\*\**p*<0.001 versus controls.

These results indicate that repeated inhalation exposure to high concentrations of CWAAP-A cause pulmonary alveolar but not bronchiolar damage in the rat. In rats exposed to CWAAP-A at 40 mg/m^3^, there was a multifocal pattern of alveolar lesions that developed into interstitial alveolar lesions with collagen deposition.

### Continuous activation of TGF**β** signaling in AEC2 by CWAAP exposure contributes to the progression of rat pulmonary disorders

TGFβ signaling is known to contribute to the pathogenesis and progression of fibrosis across multiple organs [6]. The results shown in the previous section indicate that inhalation exposure to CWAAP results in alveolitis with fibrous thickening of the interstitium. Therefore, we focused on TGFβ signaling to investigate the mechanism of lung lesions of CWAAP-A in rats. The TGFβ1 and TGFβ2 levels in the BALF are shown in Fig. 7A and 7B, respectively. Immediately after exposure and after the 2-week recovery period, both TGFβ ligands showed a significant concentration-dependent increase in the exposed group compared to the control group. After the 18-week recovery period, the induced levels of TGFβ had dropped considerably compared to the immediate post-exposure period, however, the 40 mg/m^3^ group still showed a small but significant increase in both TGFβ1 and TGFβ2 levels compared to the control group.

**Figure 7.**
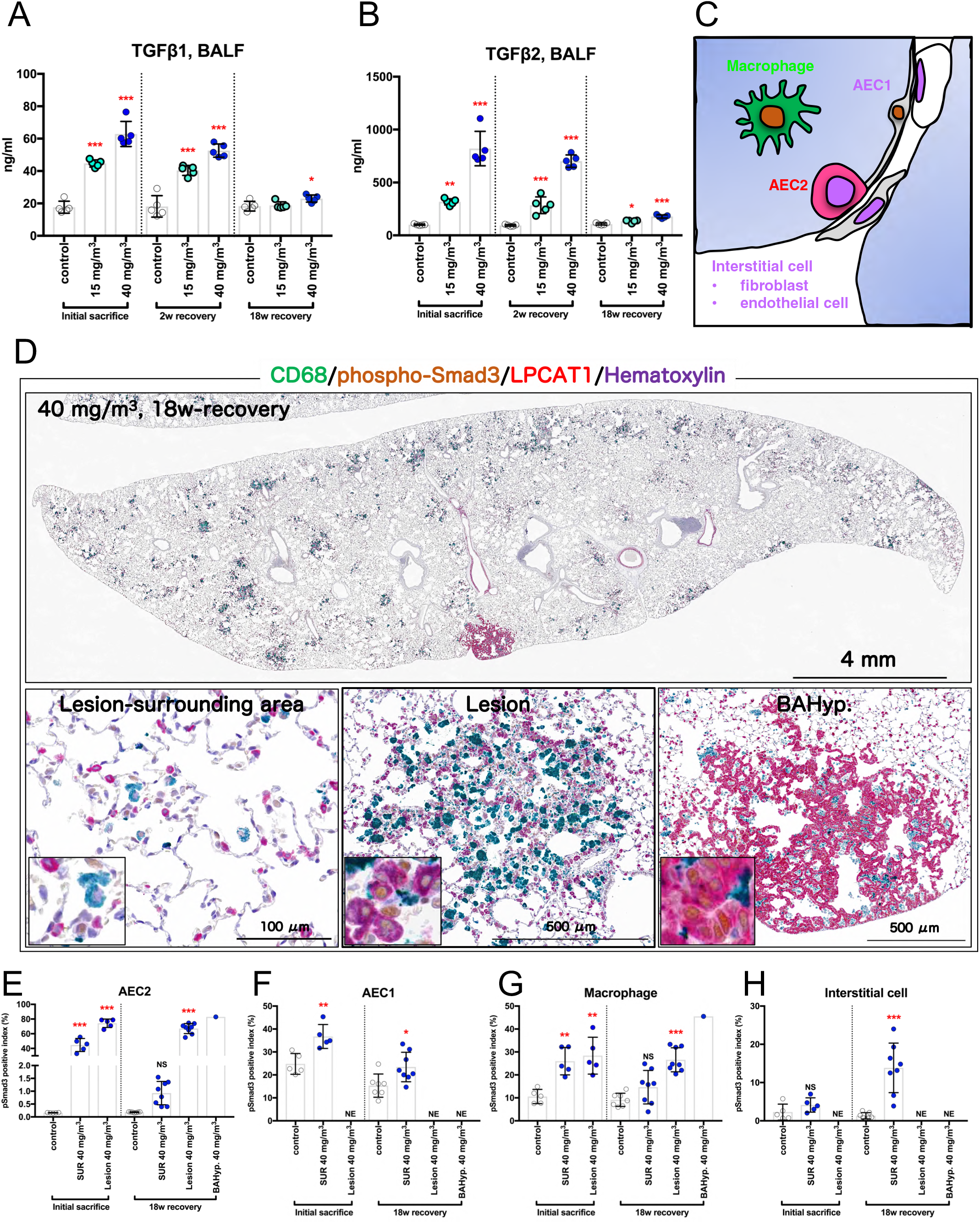
Examination for transforming growth factor (TGF) β signaling in the lung. The level of TGFβ1 (A) and TGFβ2 (B) in the BALF are shown. Schematic diagram of the alveoli after CD68-phospho-Smad3-lysophosphatidylcholine acyltransferase 1 (LPCAT1) triple staining (C): macrophages and alveolar epithelial type 2 cells (AEC2) were stained green and red, respectively, and alveolar epithelial type 1 cells (AEC1) and interstitial cells, including both fibroblast and endothelial cells, were not stained. Representative immunohistochemical staining images are shown in panel D. Nuclear phospho-Smad3 positive indexes of AEC2 (E), AEC1 (F), macrophage (G) and interstitial cell (H) are shown as bar graphs. Statistical significance was analyzed using Dunnett’s multiple comparison test: \*\**p*<0.01 and \*\*\**p*<0.001 versus controls. Abbreviations: NE, not examined; NS, not significant; SUR, normal surrounding area; and BAHyp, bronchiolo-alveolar hyperplasia.

Immunostaining with antibodies that recognize Smad3 phosphorylated at serine residues S423 and S425, lysophosphatidylcholine acyltransferase 1 (LPCAT1; an alveolar epithelial type 2 cell (AEC2) marker), and CD68 (ED-1; a macrophage marker), was used to identify the cell types in which TGFβ signaling was activated. The TGFβ signaling responder cells show brown nuclei (phospho-Smad3-positive), macrophages show green cytoplasm (CD68-positive), and AEC2 cells show red cytoplasm (LPCAT1-positive) (Fig. 7C). The alveolar epithelial type 1 cell (AEC1) has a nucleus protruding into the alveolar space, and alveolar interstitial cells are mainly vascular endothelial cells and fibroblasts in the alveolar interstitium, which are all CD68-LPCAT1 double negative (Fig. 7C). AEC2 was mainly visualized in bronchiolo-alveolar hyperplasia, while AEC2 and macrophages were mixed in multifocal lesions in agreement with the pathological morphology (Fig. 7D). In normal alveolar tissue, the cell type with the lowest phospho-Smad3 positivity was AEC2 (Fig. 7E). However, the nuclear expression of phospho-Smad3 was most markedly increased in AEC2 immediately after the end of the CWAAP-A exposure period. This high level of positive cells was seen in both the lesions and in the surrounding tissues (Fig. 7E-H), with the positive index in the lesion reaching around 80% (Fig. 7E). Interestingly, after the 18-week recovery period, the foci in the lungs of rats in the 40 mg/m^3^ group continued to show a marked increase of phospho-Smad3 positivity in AEC2, while a significant increase in the surrounding tissues disappeared (Fig. 7E). Furthermore, AEC2 in bronchiolo-alveolar hyperplasia (BAHyp) and AEC2 in multifocal lesions were found to be highly positive for phospho-Smad3 (Fig. 7D-H). These results indicate that CWAAP-A increases TGFβ ligands in the lung and that TGFβ signaling is markedly elevated, especially in the AEC2 cells in CWAAP-A induced lesions. Furthermore, data after the 18-week recovery period showed that TGFβ signaling was still markedly elevated in AEC2 cells within the lesion despite a marked decrease in lung TGFβ ligand levels. The continuous activation of TGFβ signaling in AEC2 cells within the lesion may play an essential role in the progression to fibrotic interstitial lesions.

### Alveolar epithelial progenitor cells (AEPs) are conserved in the rat lung, and expansion of AEPs are responsible for the progression of CWAAP-induced pulmonary disorders

As described above, in rats exposed to CWAAP-A, alveolar lesions centered on AEC2 were consistently observed from immediately after exposure to the end of the 18-week recovery period. Recently, it has been reported that AEPs exist in mice and humans, and play an important role in alveolar regeneration [7, 8]. We investigated the hypothesis that AEPs may be involved in CWAAP-induced rat lung lesions. Based on the report by Zacharias et al. that transmembrane 4 L six family member 1 (Tm4sf1) is an AEP marker [10], we performed immunohistochemistry for Tm4sf1. We found that alveolar epithelium highly expressing Tm4sf1 was only present in CWAAP-induced lung lesions (Fig. S5, middle panels). Interestingly, the expression of Tm4sf1 in bronchiolo-alveolar hyperplasia is as low as that in the surrounding tissues (Fig. S5, left panel, right panel), suggesting specificity of high Tm4sf1 expression in CWAAP-induced lung lesions. Double staining for LPCAT1 (an AEC2 marker) and Tm4sf1 (Fig. 8A) revealed that LPCAT1 positive cells with high expression of Tm4sf1, which identifies Tm4sf1^high^AEC2 cells as AEP cells, were consistently observed in the lesions from immediately after exposure to the end of the 18-week recovery period (Fig. 8B). However, AEPs (Tm4sf1^high^AEC2) were not observed in normal lungs or bronchiolo-alveolar hyperplasia (Fig. 8B).

**Figure 8.**
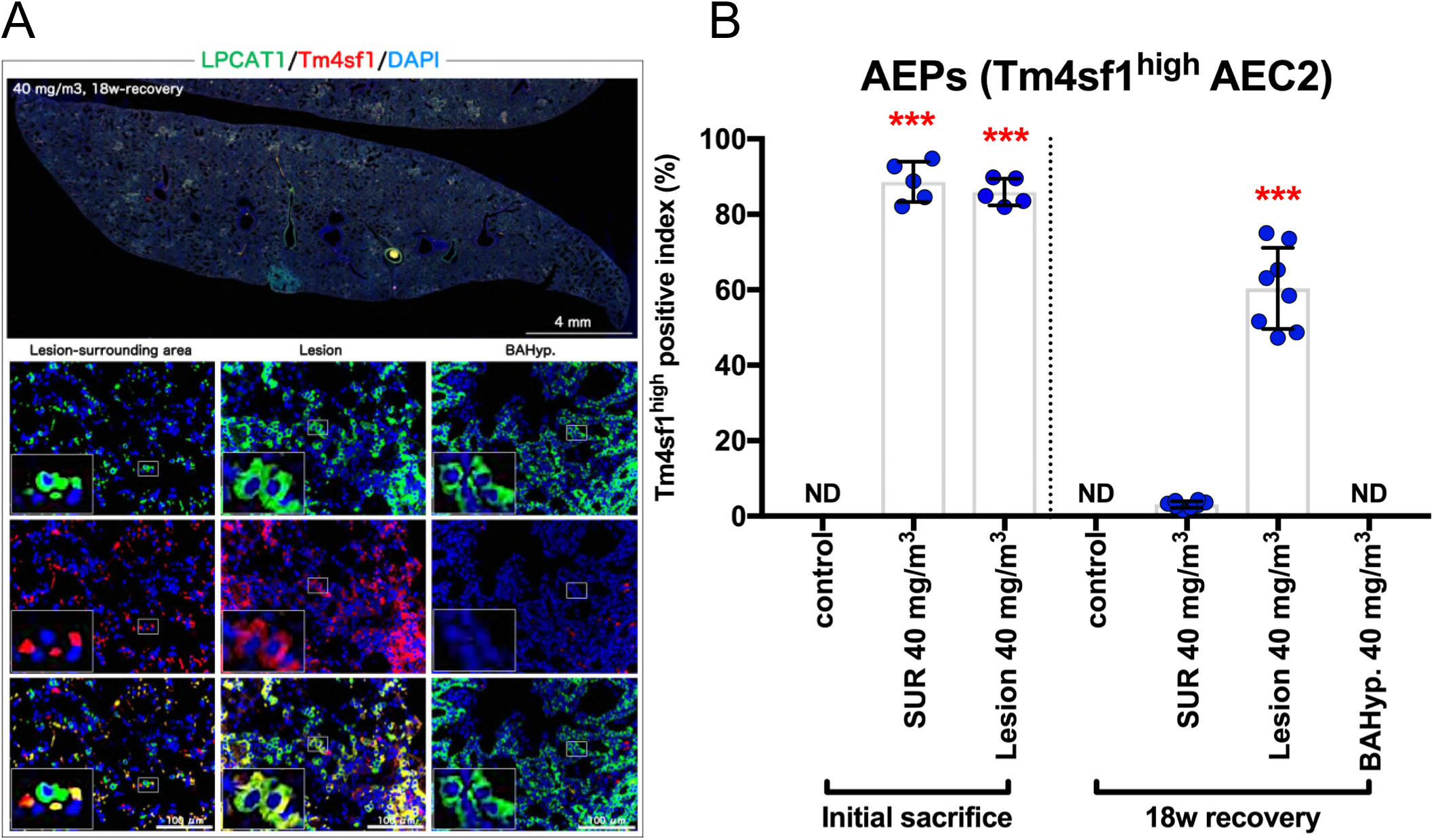
Examination of alveolar epithelial progenitor cells (AEPs) in the lung. Representative images of LPCAT1 (an AEC2 marker, green), transmembrane 4 L six family member 1 (Tm4sf1) (an AEP marker, red), and 4’,6-diamino-2-phenylindole (DAPI) (a nucleus marker, blue) co-staining (A). The Tm4sf1-high positive index was measured (B). Statistical significance was analyzed using Dunnett’s multiple comparison test: \*\*\**p*<0.001 versus controls. Abbreviation: ND, not detectable.

Laughney et al. performed single-cell analysis using various lung tumor samples and revealed the existence of Sox9-positive AEP cells as a variant AEP [11]. We then examined whether Sox9-positive AEPs are present in CWAAP-induced lung lesions. To investigate the usefulness of RT2-70 antibody as a cell membrane marker of AEC2 [12, 13], RT2-70 was double-stained with LPCAT1 or ATP binding cassette subfamily A member 3 (ABCA3), which are cytoplasmic markers of AEC2. RT2-70 was co-expressed in almost all ABCA3-positive and LPCAT1-positive cells in both normal and CWAAP-exposed lung (Fig. S6), thus confirming RT2-70-positive cells are AEC2s.

The result of triple-staining for Sox9, Tm4sf1 and RT2-70 showed that Sox9-positive AEC2s and AEPs (Tm4sf1^high^AEC2) were not observed in normal lungs (Fig. 9 left panels), while a few Sox9-Tm4sf1-RT2-70 triple-positive cells were observed in 40 mg/m^3^ exposed lungs after the 18-week recovery period (Fig. 9 right panels).

**Figure 9.**
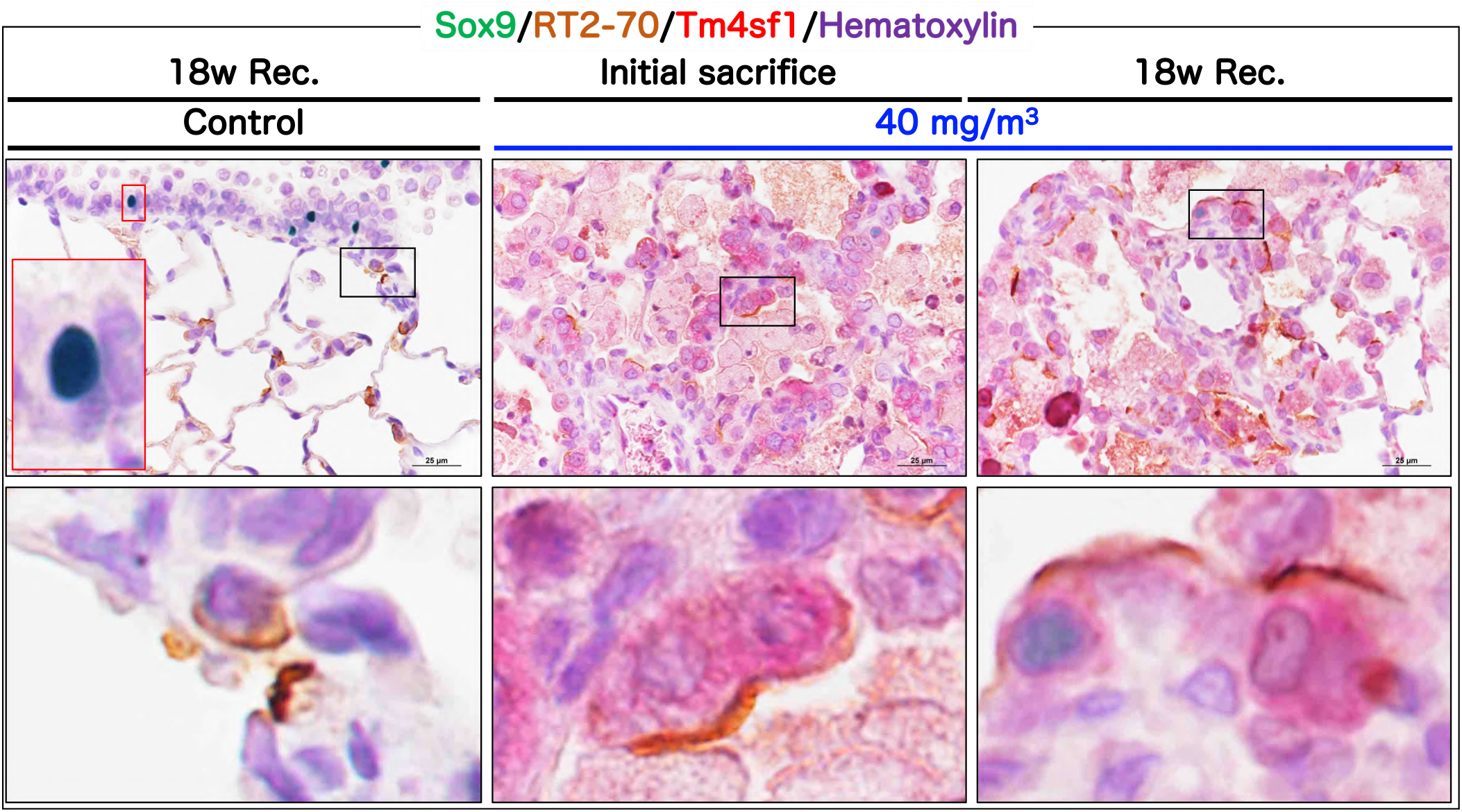
Triple staining for Sox9 (green in nucleus), RT2-70 (an AEC2 marker, brown in cell membrane), and Tm4sf1 (an AEP marker, red in cytoplasm) in the rat lung.

These results indicate that AEPs are also conserved in the rat lung, and may be involved in the formation and progression of CWAAP-induced lung lesions. Furthermore, these results support the premise that AEP expansion in the lesion represents a regenerative change in the alveoli in response to alveolar toxicity caused by exposure to CWAAP.

### CWAAP causes alveolar injury in the acute phase

In order to clarify what happened in the early stage of alveolar lesion development induced by CWAAP-A, a single inhalation exposure test was conducted. In accordance with the repeated inhalation exposure study, a large number of neutrophils and CWAAP were found in the BALF 3 days after exposure (Fig. 10A). Furthermore, the LDH activity in the BALF showed a significant increase compared to the control group at 1 hour and 3 days after exposure in both the 40 and 100 mg/m^3^ groups (Fig. 10B). The significant increase in the total number of cells in the BALF in the exposure groups three days after exposure was clearly due to neutrophils (Fig. 10C, E). These results indicate that a single exposure of CWAAP causes significant increases in LDH activity in the BALF and induces neutrophil infiltration into the lungs. To obtain more accurate pathological images of the lungs, we developed a method to inflate lungs with air only, without injecting formalin solution into the lungs which is routinely done (Fig. S7). In this method, the trachea is clipped before opening the chest to prevent air from escaping from the lung exposed to atmospheric pressure (Fig. S7A). Histopathological images of the left lung inflated with air alone and the right lung inflated with formalin injection were compared, and good expansion of alveoli was confirmed in both cases (Fig. S7B). Histopathological evaluation of the left lung of the single-exposed animals using this method revealed a large number of neutrophils around the collapsed alveoli in the alveolar region (Fig. 11A), and the surrounding alveolar ducts showed overinflation (Fig. 11A center and right panels). These pathological findings are shown in Table 2.

**Figure 10.**
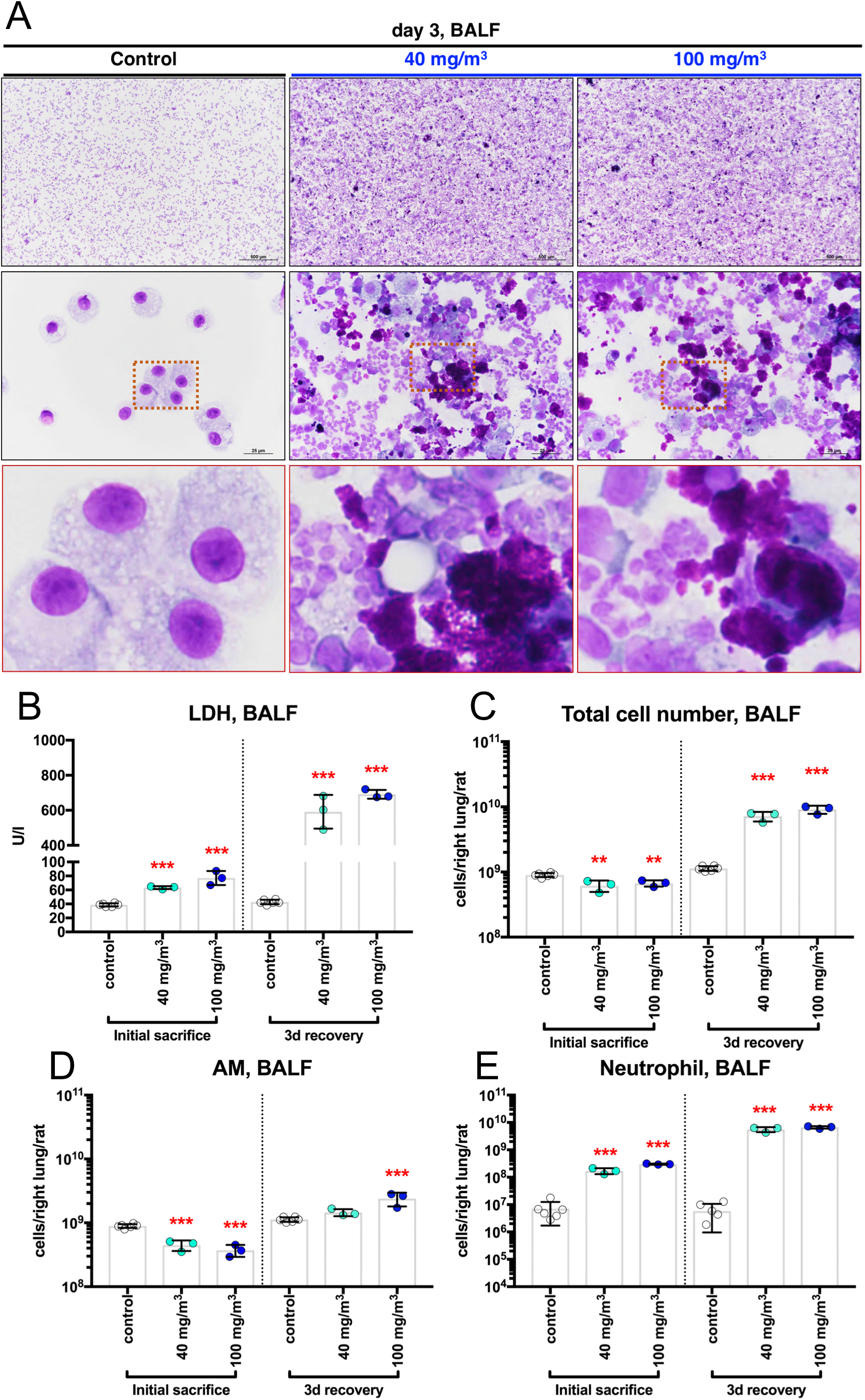
BALF was collected from male rats after a single inhalation exposure to CWAAP-A (40 or 100 mg/m^3^). Representative images of the BALF cytospin cytology (A). The lactate dehydrogenase (LDH) activity (B), total cell number (C), alveolar macrophage number (D), and neutrophil number (E) in the BALF. Statistical significance was analyzed using Dunnett’s multiple comparison test: \*\**p*<0.01 and \*\*\**p*<0.001 versus controls. Abbreviation: 3d, day 3

**Figure 11.**
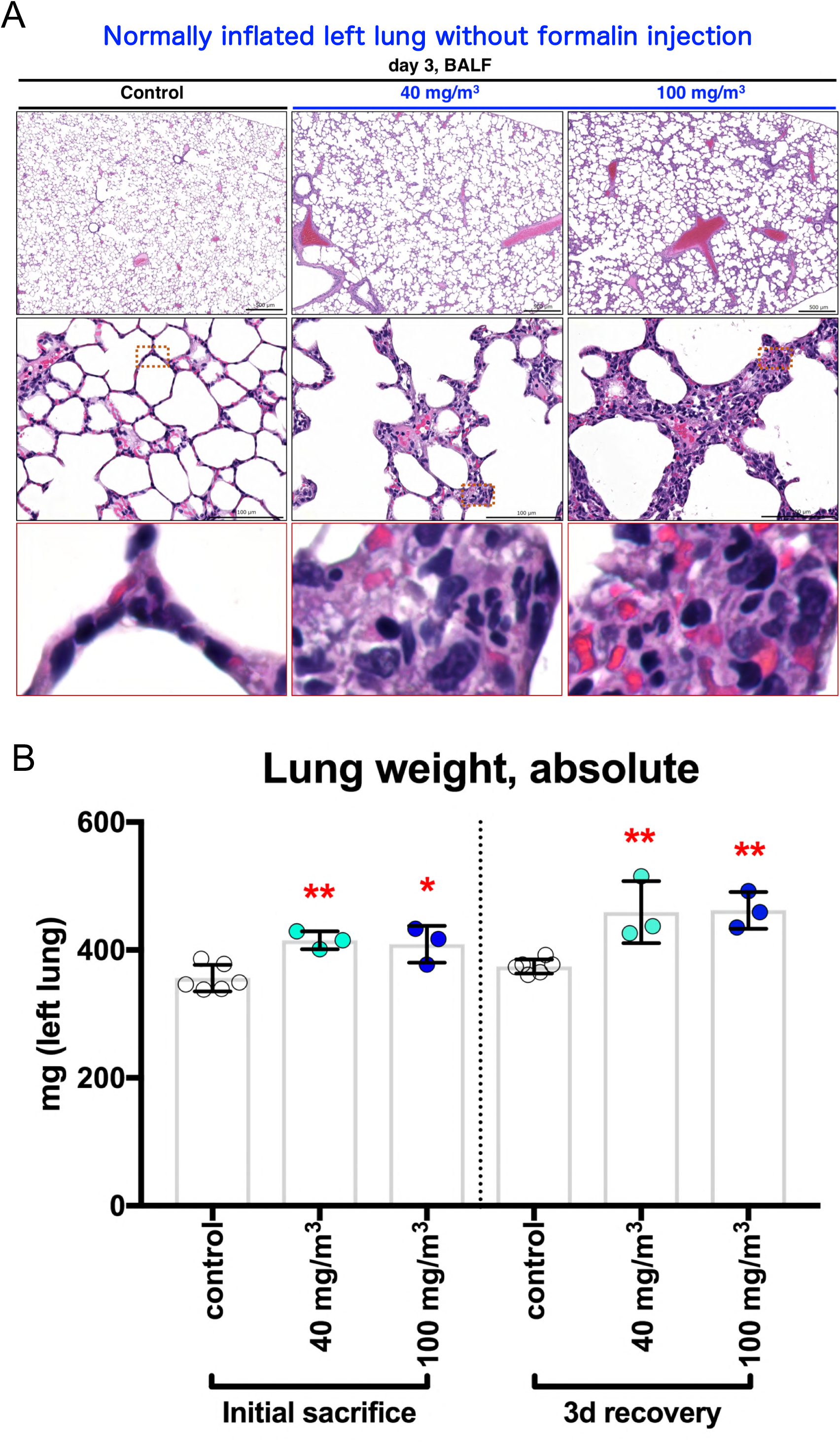
The effect of a single inhalation exposure to CWAAP-A on the pathology and weight of the lungs. Representative histopathological photographs of the left lung inflated by air (A) and left lung weight (B). Statistical significance was analyzed using Dunnett’s multiple comparison test: \**p*<0.05 and \*\**p*<0.01 versus controls.

**Table 2.**
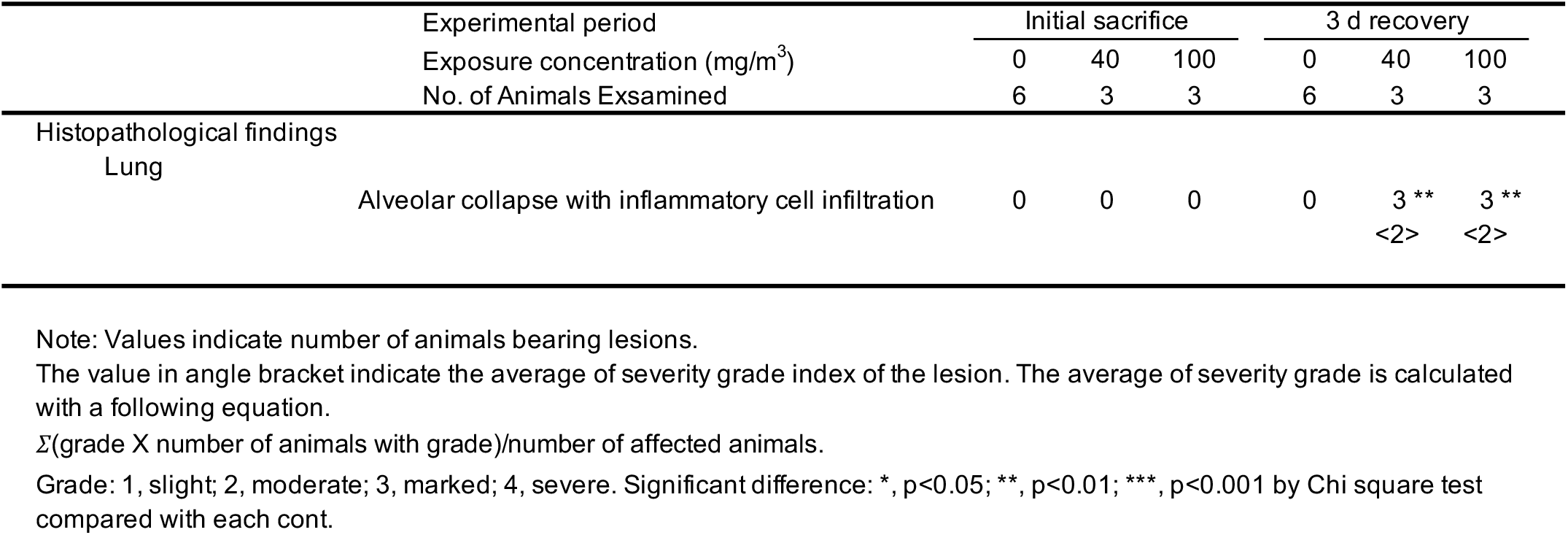
Histopathological findings of the lung after a single inhalation exposure to CWAAP-A.

Alveolar collapse with inflammatory cell infiltration was specifically observed animals 3 days after exposure to 40 and 100 mg/m^3^ CWAAP-A. Interestingly, these findings were not observed in the pathological specimens of the right lung, which were inflated by formalin injection. (Fig. S8). In addition, a significant increase in lung weight was observed in both the 1-hour and 3-day post-exposure autopsy groups (Fig. 11B). These results indicate that a single inhalation exposure to CWAAP-A causes alveolar injury characterized by alveolar collapse with a high degree of neutrophilic infiltration in the acute phase.

### Intratracheal instillation is useful for screening of CWAAPs-induced rat pulmonary disorders

Although the intratracheal instillation method is useful for low-cost and simple toxicity assessment of a large number of chemicals, it is unclear whether this can be applied to CWAAPs, which are characterized by very high water-absorption. Therefore, we conducted an intratracheal instillation study using CWAAP-A, which was used in the whole-body inhalation exposure study, and another CWAAP, CWAAP-B. The administration and sampling schedule was similar to that of the inhalation exposure study. The final body weight was not affected by the intratracheal administration of either of the CWAAPs (Fig. 12A). In sharp contrast, lung weight (Fig. 12B) and mediastinal lymph node weight (Fig. 12C) were dramatically increased in the 2-week recovery group. The increases in lung weight and mediastinal lymph node weight were less after the 18-week recovery period, but they were still significantly elevated. A similar trend was also observed for other lesion markers: total cell count in the BALF (Fig. 12D), alveolar macrophage count (Fig. 12E), neutrophil count (Fig. 12F), LDH activity (Fig. 12G), SP-D level (Fig. 12H) in the BALF, and SP-D level in the plasma (Fig. 12I) were all markedly increased compared to controls 2 weeks after CWAAP administration and these increases had decreased or disappeared at 18 weeks after CWAAP administration. Representative lung and histopathological images are shown in Fig. 13. Because CWAAPs were administered intratracheally as a liquid suspension in phosphate-buffered saline (PBS), the foci were distributed caudally in the left and right lungs (Fig. 13). The histopathological findings of the lungs and mediastinal lymph nodes observed in this study are shown in Table 3 and are qualitatively similar to the results of the systemic inhalation exposure study (Table 1).

**Figure 12.**
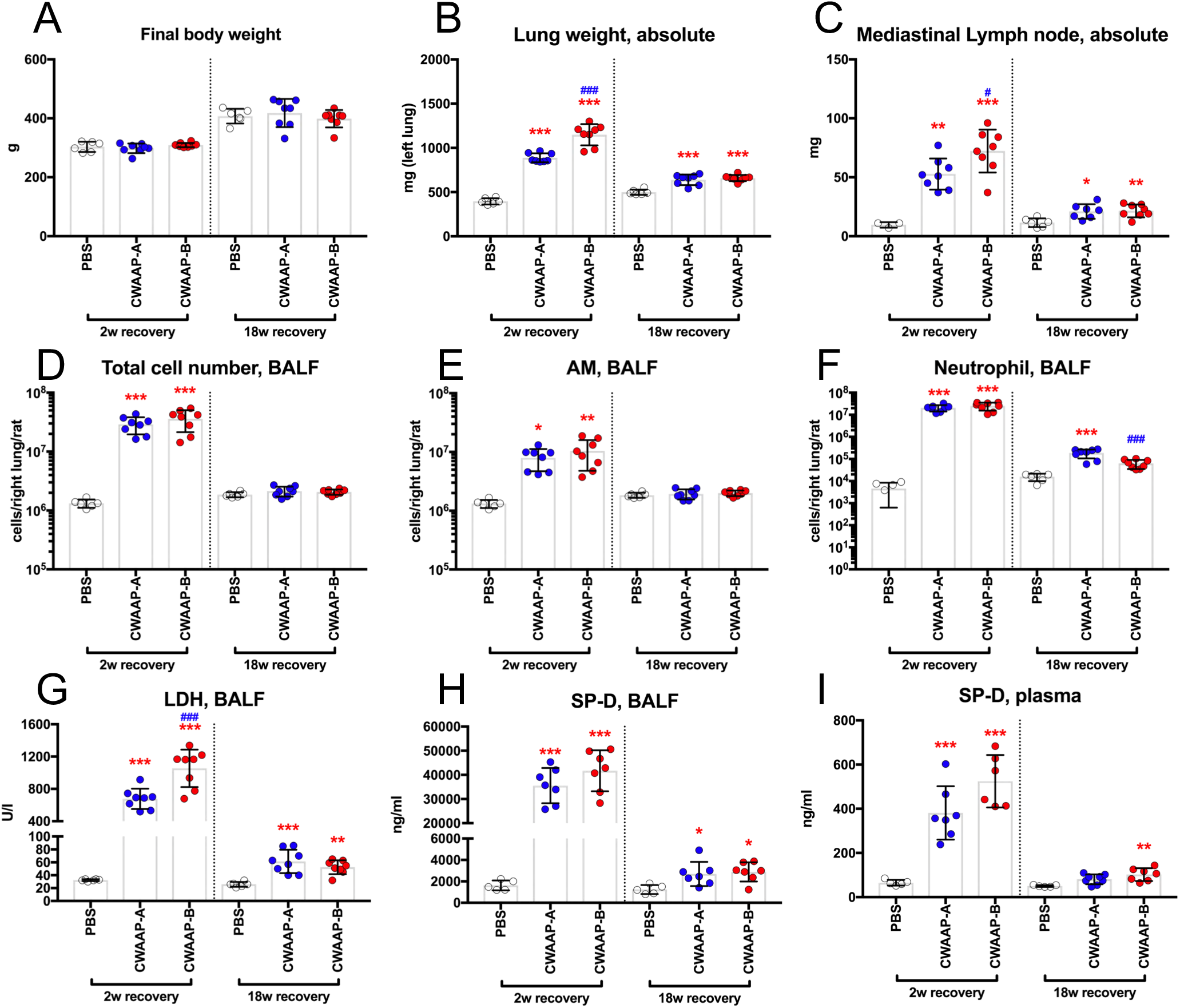
CWAAP-A or CWAAP-B (both administered at 1 mg/kg per dose) were administered to male rats by repeated intratracheal instillations (see Fig. S10C for details). Final body weights (A), left lung weights (B), mediastinal lymph node weights (C), total cell number in the BALF (D), alveolar macrophage number in the BALF (E), neutrophil number in the BALF (F), LDH activity in the BALF (G), SP-D level in the BALF (H), and SP-D level in the plasma (I) were measured at each sacrifice. Tukey’s multiple comparison tests were carried out for each sacrifice: \**p*<0.05, \*\**p*<0.01, and \*\*\**p*<0.001 indicate significant differences from the control (PBS) group, and ^#^*p*<0.05 and ^###^*p*<0.001 indicate significant differences between CWAAP-A and CWAAP-B.

**Figure 13.**
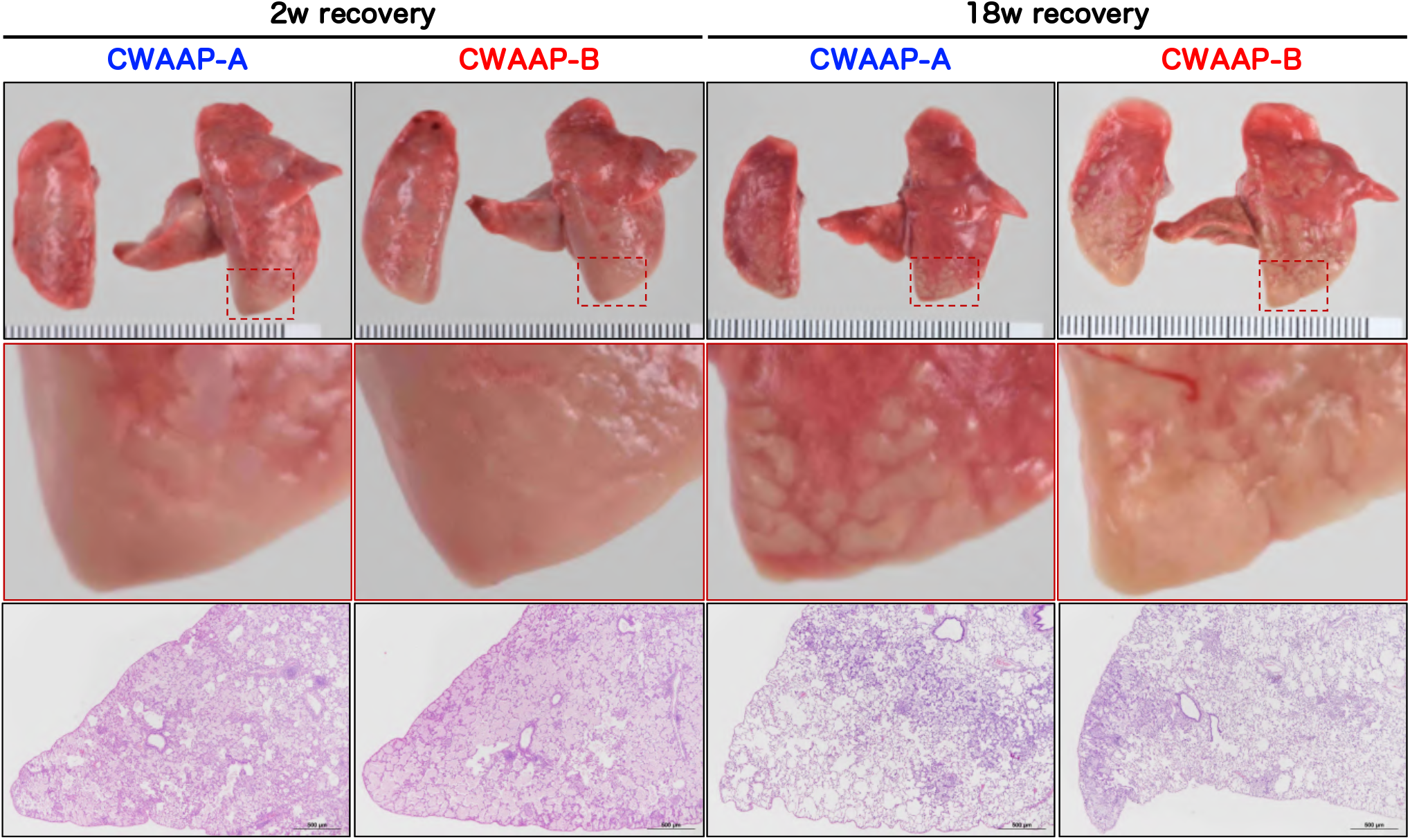
Representative macroscopic and microscopic photographs of rat lungs after intratracheal instillation of CWAAPs. White spots were observed on the caudal side of the right and left lungs (middle panels), and histopathological images of the same areas showed diffuse lesions (lower panels).

**Table 3.**
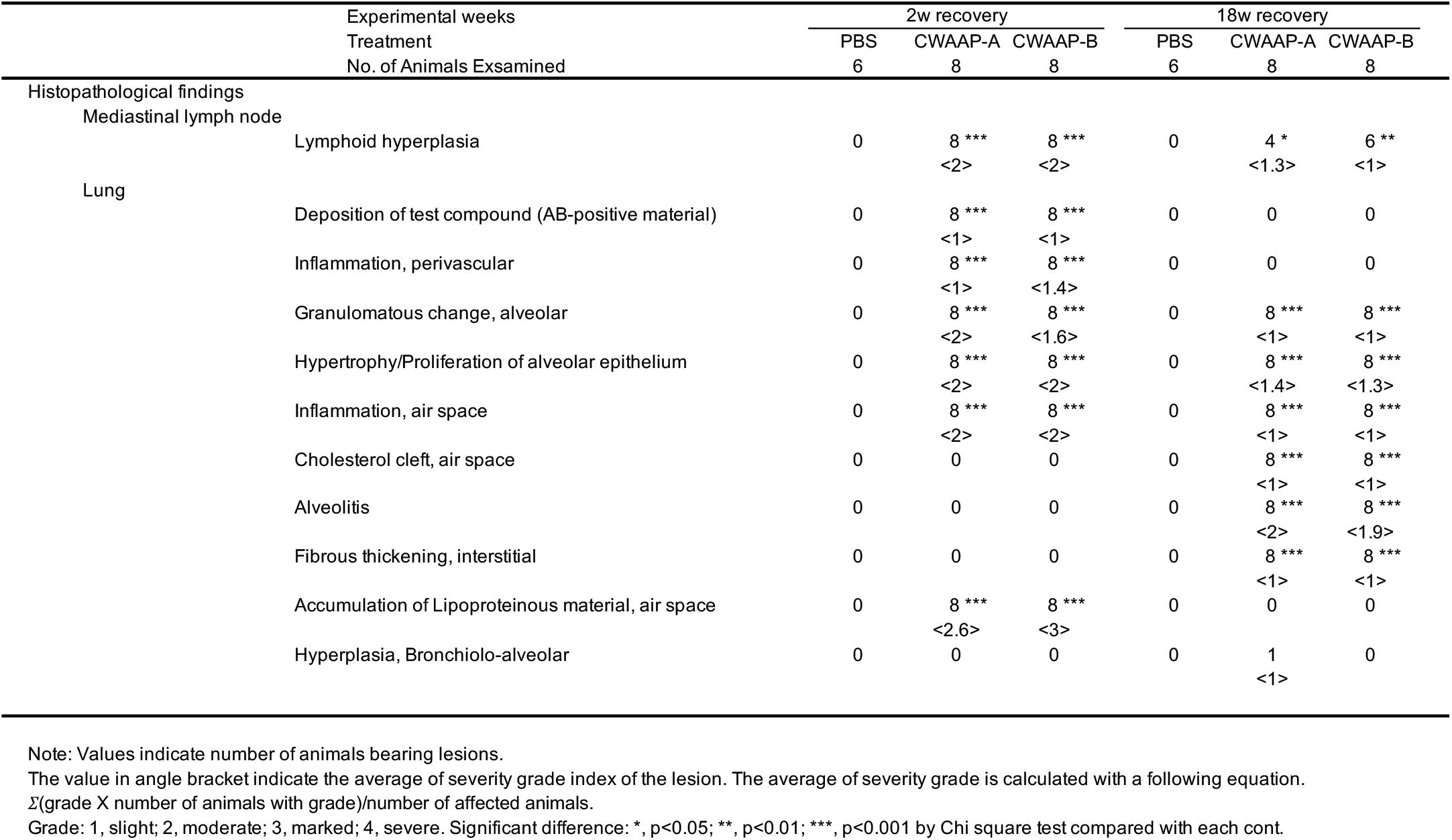
Histopathological findings of the mediastinal lymph node and lung after repeated intratracheal instillation of CWAAPs.

There were differences in the toxicity of CWAAP-A and CWAAP-B. Significant increases were observed in lung weight (Fig. 12B), mediastinal lymph node weight (Fig. 12C), and LDH level in the BALF (Fig. 12G) of rats administered CWAAP-B compared to rats administered CWAAP-A after the 2-week recovery period. Consistent with these results, the histopathological analysis showed that the accumulation of lipoproteinous material, an important indicator of alveolar proteinosis, tended to be greater in CWAAP-B than in CWAAP-A (Table 3). However, these differences disappeared after the 18-week recovery period. These results indicate that serial intratracheal administration can detect the pulmonary toxicity of CWAAPs in rats. Furthermore, it is useful for comparing the intensity of pulmonary damage caused by different CWAAPs. Our results suggest that CWAAP-B has a stronger damaging effect in the lung than CWAAP-A.

## Discussion

Kishimoto et al. in our research group recently reported detailed data on clinical pulmonary pathogenesis in workers who had handled CWAAPs, a type of organic compound [2]. Previously, exposure to CWAAPs was not considered harmful to humans. The present study reports for the first time the results of rat toxicity tests using systemic inhalation exposure and intratracheal instillation of CWAAPs used in the factory where an industrial accident occurred (see Background). We found that inhalation exposure to CWAAP-A caused alveolar collapse and neutrophil infiltration into the lung and this progressed to fibrosis. At the cellular level, TGFβ signaling in alveolar epithelial type 2 cells (AEC2) and alveolar epithelial progenitor cells (AEPs) and expansion of AEPs were primary events in the progression of CWAAP-induced pulmonary lesions. A schematic diagram of the mechanism of CWAAP-induced lung disease in rats revealed in this study is shown in Fig. 14.

**Figure 14.**
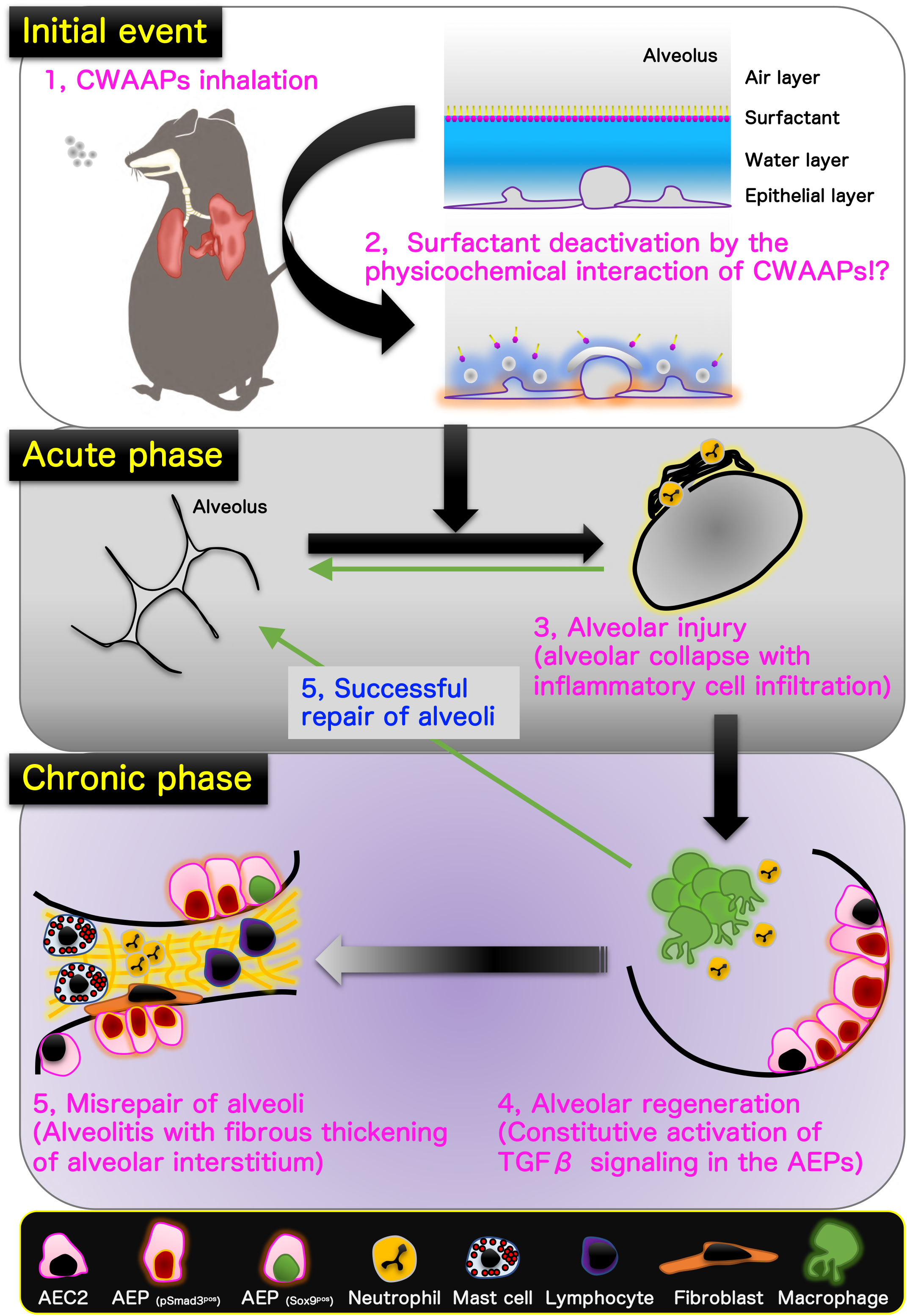
Graphical abstract for this study.

In this study, we used a single inhalation exposure to CWAAP-A to investigate the lung damage of CWAAP-A in the acute phase, and utilized an alternative sampling method (Fig. S7) that reproduces as closely as possible the inflation of rat lungs without formalin inflation. We found that in the rat lung in the acute phase after CWAAP inhalation, alveolar collapse with a high degree of neutrophil infiltration occurred (Fig. 14, middle panel). If the lung remains collapsed and the resting tissue is filled with edema, gas exchange will be defective and pneumonia will likely occur [14, 15]. If the alveoli are not reopened, the pathologic lung will progress to fibrosis [16–18]. Therefore, the CWAAP-induced alveolar collapse in the acute phase may trigger pneumonia and the lung fibrotic changes observed in the chronic phase.

Typical diseases that cause alveolar injury are acute respiratory distress syndrome (ARDS) and ventilator-induced lung injury secondary to mechanical ventilation, which is the only treatment for ARDS [19]. ARDS is a complex syndrome characterized by four well-accepted central components [20] known as the pathologic tetrad [21]. When loss of surfactant function (surfactant deactivation) [22] occurs, the surface tension of the alveoli increases, which is known to exacerbate the increase in edema fluid in the alveoli [23]. Surfactant dysfunction alters alveolar mechanics, resulting in recruitment/derecruitment of alveoli with each breath [24]. Surfactant deactivation exacerbates alveolar collapse over time by promoting instability and collapse of the heterogeneous lung tissue. Consistent with these components, the present study found that a single exposure to CWAAP resulted in alveolar collapse with neutrophilic infiltration and overinflation of alveolar ducts around collapsed alveoli in rat lungs 3 days after exposure. However, hyaline membrane formation, hemorrhage, and plasma pulmonary edema were not observed. These results suggest that CWAAP exposure induces alveolar collapse in rat lungs, but the direct damage to epithelial cells and endothelial cells is minor. It is commonly understood that CWAAPs are dispersed in water at the molecular level, and unlike SAPs swell by incorporating water into their cross-linked molecular structure [25–27]. Taken together, inhaled CWAAP may absorb water molecules on the alveolar epithelial surface (the water layer) and damage the surfactant layer, thereby altering alveolar mechanics and causing alveolar collapse and inflammation (upper panel of Fig. 14). Thus, in contrast to ARDS, which is caused by increased vascular permeability and severe epithelial/endothelial injury, the acute phase of CWAAP-induced lung lesions in rats may be due to alveolar damage secondary to the disruption of the alveolar surfactant microenvironment that occurs after CWAAP disrupts the water layer.

The present study found that alveolitis with fibrous thickening of the alveolar septa occurs as a typical chronic lesion caused by CWAAP. This lesion partially progressed as a pathological condition even though most of the initial lesions were restored to normal (alveolar regeneration, Fig. 14, lower right) after an 18-week recovery period after the final exposure to CWAAP-A (Fig. 14, lower left). Alveolitis is the basic pathological term for interstitial pneumonia. Idiopathic pulmonary fibrosis, a classic example of interstitial pneumonia with fibrosis, is one of the most studied areas of respiratory diseases [28–30], and TGFβ signaling plays a central role in the pathogenesis of pulmonary fibrotic disease [6, 31]. TGFβ signaling is activated by the binding of ligands, including TGFβ1 and TGFβ2, to their receptors on the cell surface, and regulates downstream transcriptional networks through phosphorylation and nuclear translocation of Smad2 and Smad3 [7]. Phosphorylation of serine residues (S423 and S425) at the C-terminus of the MH2 domain of Smad3 protein is essential for activating the transcriptional network by TGFβ signaling [32]. In this study, using an antibody that recognizes phosphorylated serine residues of Smad3, we found that persistent TGFβ signaling was activated in AEC2s in multifocal lesions even after an 18-week recovery period following CWAAP-A exposure. A previous study using a mouse model of lipopolysaccharide-induced lung injury reported that activation of TGFβ signaling in AEC2s is required for cell cycle arrest but inhibits differentiation to AEC1 [33]. Further studies focusing on the molecular mechanisms by which TGFβ signaling in proliferating AEC2s continues to be activated are needed to better understand the pulmonary toxicity of CWAAPs.

CWAAP-induced alveolar lesions consist of enlarged and proliferating AEC2s for alveolar regeneration. Recently, single-cell analysis has become widely used to identify the cell populations that compose normal and diseased lungs, especially disease-specific cell populations [34–36]. It has been reported that AEPs are present in mouse and human lungs to contribute to the activation of regenerative molecular programs [9, 10] and that Sox9-positive AEPs also exist [11]. Although in this study, we were not able to identify AEPs in normal rat lungs, AEPs were observed immediately after CWAAP exposure and after the 18-week recovery period, suggesting that CWAAP-induced lung lesions are regenerative changes, consistent with previous reports. Furthermore, a small number of Sox9-positive AEPs were also observed in the lesions after an 18-week recovery period. This study is the first report to discuss the presence of AEPs in rat lung lesions.

This study did not identify the mechanism of alveolitis with fibrous thickening of the alveolar interstitium induced by CWAAP. It is well known that fibroblasts transdifferentiated from AEC2s via epithelial-mesenchymal transition (EMT) are candidates for collagen-producing cells in pulmonary fibrosis [31, 37]. TGFβ signaling is also an important signal for the induction of EMT [6]. Recently, Li et al. reported that the TGFβ-Sox9 axis produces collagen 10a1 in EMT-mediated gastric cancer cells, and demonstrated that Sox9 binds directly to the promoter region of the col10a1 gene [38]. We found a high ratio of both phospho-Smad3-positive AEC2s and AEPs within CWAAP-induced lung lesions. Since CWAAP-induced lung lesions transitioned to alveolar interstitial-predominant lesions in the chronic phase, an EMT-mediated trans-differentiation of AEPs into fibroblast and involvement of reprograming via Sox9 expression may have occurred. These possibilities need to be studied in the future.

The present study is the first to investigate the pulmonary toxicity of CWAAP in rats by both inhalation exposure and intratracheal instillation. The results showed that the pathological lesions caused by intratracheally administered CWAAP-A were qualitatively similar to those caused by inhalation exposure, and changes in plasma and BALF biochemical and cytological parameters were also similar in both studies. These findings indicate that intratracheal instillation is useful for the initial assessment of acute and chronic toxicity of CWAAPs. Furthermore, using intratracheal instillation a second CWAAP, CWAAP-B, was evaluated. Two weeks after the final administration of CWAAPs there was a significant increase in lung weight and higher LDH activity in the BALF of the rats administered CWAAP-B compared to CWAAP-A, suggesting that CWAAP-B may have higher pulmonary toxicity than CWAAP-A. Thus, intratracheal instillation, which is rapid, simple, and inexpensive, is effective for the initial investigation of pulmonary toxicity of different CWAAPs.

The present study has two limitations. First, while we succeeded in clarifying the pulmonary toxicity of CWAAP-A in rats in the acute phase up to the chronic phase, since all animal experiments were evaluated only up to 26 week we could not clarify the end-stage pathogenesis of CWAAP-treated lungs, such as whether alveolitis with fibrous thickening progresses to pulmonary fibrosis. This may be the reason that in this study there was no evidence of progressive fibrosis with a high degree of destruction and remodeling of alveolar structures. Notably, one case of bronchiolo-alveolar hyperplasia, a pre-neoplastic lesion, was observed in one CWAAP-A-exposed rat in the inhalation study and one CWAAP-A treated rat in the intratracheal instillation study. To evaluate any lung carcinogenicity associated with inflammatory-fibrotic diseases caused by CWAAP, longer-term studies are needed. Second, we adopted an irregular inhalation exposure protocol of four hours a day, once a week at high concentrations, assuming the conditions of an exposure site where an industrial accident occurred. In addition, to ensure a sufficient number of animals for each condition, only two concentrations were considered for the study. To clarify the details of the dose-response relationship and recovery from CWAAP, it is necessary to conduct a whole-body inhalation exposure study using an exposure protocol of 6 hours per day, 5 days per week at more than 3 concentrations in compliance with the test guideline (TG413) by the Organization for Economic Co-operation and Development [39]. We are currently conducting another study using the above protocol and are in the process of summarizing the results.

## Conclusions

In this study, we investigated the pathological/toxicological characteristics of CWAAP, which has caused occupational lung diseases in Japanese workplaces in recent years. The results showed that exposure to CWAAP caused alveolar injury in the acute phase and that repeated exposures caused regenerative changes in the alveoli. After the end of exposure, during the recovery period, the alveolar lesions partially recovered to normal, but the some progressed to alveolitis with fibrous thickening. TGFβ signaling in AEC2 and AEP cells and AEP cell expansion were primary events in the pathogenesis of CWAAP. Moreover, we compared the lung pathology of CWAAP administered by systemic inhalation exposure with that of CWAAP administered by intratracheal instillation and found that the lung pathologies were similar. The use of intratracheal instillation as an adjunct to inhalation exposure is expected to greatly accelerate testing of respirable CWAAP and SAP products, and consequently, to significantly increase our understanding of the pulmonary toxicity of CWAAP and SAP products.

## Methods

### Materials

CWAAP-A and CWAAP-B were purchased from a company which produces various polymers. As shown in Fig. S3, CWAAP-A seems to be finer and lighter than CWAAP-B. A list of all primary antibodies used in these studies is summarized in Table S2. The donkey anti-mouse IgG conjugated Alexa Fluor 488 (ab150105) and anti-rabbit IgG conjugated Alexa Fluor 594 (ab150064) were purchased from Abcam plc (Cambridge, UK). The VECTASHIELD Mounting Medium with DAPI (H-1200) was purchased from Vector laboratories (Burlingame, CA). The other reagents were of the highest grade commercially available.

### Animals

All animal experiments were approved by the Animal Experiment Committee of the Japan Bioassay Research Center. Male F344 rats were purchased from Charles River Laboratories Japan, Inc. (Yokohama, Japan) and Japan SLC, Inc. (Hamamatsu, Japan). The rats were housed in an air-conditioned room under a 12 hours light/12 hours dark (8:00-20:00, light cycle) photoperiod, and fed a general diet (CRF-1, Oriental Yeast Co. Ltd., Tokyo, Japan) and tap water *ad libitum*. After 1-2 weeks of quarantine and acclimation, they were treated with CWAAPs.

### Whole body inhalation

Whole body inhalation was conducted using the direct-injection whole body inhalation system (Shibata Scientific Technology, Ltd., Soka, Japan) (Fig. S9A). To prevent the CWAAP from absorbing moisture, the system has been modified to supply dry air into the port of the cartridge. Rats were exposed to CWAAP-A by either repeated or single exposures (see Fig. S10A and S10B). The exposure concentrations of CWAAP-A were based on previous feasibility experiments (data not shown) and the concentration of CWAAP in a workplace where workers were exposed to CWAAPs. The workplace concentration was more than 40 mg/m^3^ [2], therefore, the target concentration of 40 mg/m^3^ was used in both the repeated exposure experiment (15 and 40 mg/m^3^) and single exposure experiment (40 and 100 mg/m^3^). In both experiments, the duration of exposure was 4 hours. For the repeated exposure experiment, rats were exposed to CWAAP-A once a week for 2 months (9 times in total). Two inhalation chambers were used for each target concentration to ensure that an adequate number of animals were exposed to each concentration of CWAAP-A (No.1 and 2: 15 mg/m^3^; and No. 3 and 4: 40 mg/m^3^). Inhalation exposure to CWAAPs was started once the rats were 11 weeks old.

Aerosol generation for CWAAP-A was conducted according to the manufacture’s instructions. CWAAP-A was weighed and placed into the inner cartridge, which was then set into the dedicated port of the inhalation system. Compressed air was injected into the cartridge to generate an aerosol, which was then fed into the inhalation chamber. Compressed air at 0.4 Mpa was injected three times with a duration of 0.2 seconds and an interval of 0.3 seconds to empty the CWAAP from the cartridge. The interval between each series of injections was 4 minutes per cartridge (i.e., 60 cartridges were required for a 4-hour exposure). The humidity (30∼40%) in the chamber was adjusted using a valve linked to the humidification bottle. Once the humidity in the inhalation chamber was below 40%, the animals were placed in the chamber and exposure was initiated.

During exposure, the number of particles in each inhalation chamber was monitored by an optical particle controller (OPC-AP-600, Shibata Scientific Technology), and the concentration of CWAAP-A in the inhalation chamber was measured at least twice from hour 1 to hour 3 after the start of exposure by collecting the test substance in the inhalation chamber on a fluoropolymer binder glass fiber filter (TX40HI20-WW, φ55 mm, Tokyo Dylec, Corp., Tokyo, Japan). The particle size of CWAAP-A in the chamber was measured using a micro-orifice uniform deposit cascade impactor (MOUDI-II, MSP Corp., Shoreview, MN) during the second and the eighth exposures during the repeated exposure experiment. The mass median aerodynamic diameter (MMAD) and geometric standard deviation (σg) were calculated by cumulative frequency distribution graphs with logarithmic probability. In addition, CWAAP-A in the inhalation chamber was collected on a 0.2 μm polycarbonate filter (φ47 mm, Whatman plc, Little Chalfont, UK) and observed by scanning electron microscope (SEM) (SU8000, Hitachi High-Tech, Tokyo, Japan). The measured concentration of CWAAP-A in the repeated exposure experiment are shown in Fig. S9B. The measured concentrations of CWAAP-A in the chamber were maintained at approximately the target concentrations throughout the exposure period. The MMADs were 0.8 μm (Fig. S9D and S9E) with σg values below 2.5 (Fig. S9E). The coefficient of variation of the No.4 chamber was high because a system error occurred due to the jamming of the cartridge in the cartridge holder during the seventh exposure. However, we confirmed that this error was resolved within one hour, and the average CWAAP concentration was maintained throughout the exposure time. The CWAAP-A particles generated in the chamber did not appear to be highly aggregated or humidified (Fig. S9C). In the single exposure experiment, the measured concentrations of CWAAP-A were 42.5 mg/m^3^ in the 40 mg/m^3^ chamber and 100.4 mg/m^3^ in the 100 mg/m^3^ chamber, σg values were similar to those of the repeated exposure experiment. These data indicate that CWAAP-A exposure was stable throughout the exposure periods in both the repeated and single exposure experiments.

In the repeated exposure experiment, rats were euthanized by exsanguination under isoflurane anesthesia immediately after the last exposure and at 2 weeks and 18 weeks after the last exposure. In the single exposure test, rats were sacrificed immediately after exposure and 3 days after exposure. BALF was collected as described below, and the lungs from which BALF was not collected were weighed and then fixed in 10% neutral phosphate-buffered formalin solution.

### Intratracheal instillation

CWAAP-A and CWAAP-B were suspended in PBS, sonicated and neutralized by 1M NaOH. The final concentration of the CWAAP solution was 1 mg/ml. Our preliminary study suggested that a single intratracheal administration of 1.5 mg/kg would have too strong an effect on the lung, while a dose of 0.5 mg/kg would be much weaker and return to normal in a few weeks (Fig. S11). In addition, a previous study reported that a single intratracheal administration of CWAAP at 0.1 mg/rat (about 0.43 mg/kg) caused a moderate inflammatory response [5]. Therefore, the dose of CWAAPs was set at 1 mg/kg, and the administration interval set at two weeks. For comparison with the whole body inhalation study, CWAAPs were administered over 2 month period (a total of 5 times) (Fig. S10C). The Z-average (median particle diameter) and zeta potential of the preparations were measured using Zetasizer Ultra (Malvern Panalytical Ltd., Worcestershire, UK): CWAAP-A, 471±136 nm and -35.7±1.7 mV; and CWAAP-B, 705±53 nm and -37.5±2.5 mV, respectively. Intratracheal instillations of CWAAPs to rats was started when the rats were 8 weeks old. Rats were placed under isoflurane inhalation anesthesia and CWAAP solution was injected at 1 mg/kg into the trachea of rats using a MicroSprayer^®^ Aerosolizer (Model IA-1B; Penn-Century, Inc., Wyndmoor, PA). At 2 weeks and 18 weeks after the final CWAAP administration, rats were placed under isoflurane anesthesia and euthanized by exsanguination. BALF was collected as described below. The left lung was then weighed and fixed in 10% neutral phosphate buffered formalin solution.

### BALF collection hand analysis

The left (intratracheal instillation study and the single inhalation study) or right (repeated inhalation study) bronchus was tied off with a thread, and the opposite lung lobes were lavaged with 4-8 ml of saline. The total cell number in the BALF was counted using an Automated Hematology Analyzer (XN-2000V; Sysmex Corp., Kobe, Japan). Cell populations on glass slides were prepared using Cytospin 4 (Thermo Fisher Scientific, Inc., Waltham, MA). After May-Grunwald-Giemsa staining, the differential white blood cell count was made by visual observation. To measure LDH activity, the BALF was centrifuged at 1,960 rpm (800 × *g*) for 10 min at 4°C, and the supernatant was examined using an automatic analyzer (Hitachi 7080 or 7180, Hitachi High-Tech Corp., Tokyo, Japan).

### Enzyme immhunoassay

SP-D concentrations in the BALF and plasma were determined using a Rat/Mouse SP-D kit YAMASA EIA (YAMASA Corp., Choshi, Japan). In this assay, the BALF was diluted 500- or 1,000-fold, and plasma was diluted 25-fold with the assay diluent included in the kit. BALF concentrations of TGFβ1 and TGFβ2 were measured by Human/Mouse/Rat/Porcine/Canine TGF-beta 1 Quantikine ELISA Kit (R&D Systems) and Mouse/Rat/Canine/Porcine TGF-beta 2 Quantikine ELISA Kit (R&D Systems). For the TGFβ2 assay, the BALF was diluted 3-fold with the sample diluent included in the kit. The absorbance at 450 nm was measured using a microplate reader (Spark^®^; Tecan Group, Ltd., Männedorf, Switzerland or SpectraMax; Molecular Devices, LLC., San Jose, CA). For TGFβ1 and TGFβ2 assays, the absorbance at 570 nm was also measured and subtracted as a background signal.

### Hydroxyproline assay

The content of hydroxyproline, a main component of collagen, in the lungs was measured using a hydroxyproline assay kit (Perchlorate-Free) (BioVision, Inc., San Francisco, CA). Small pieces of lung (about 30 mg) were homogenized in 10 volumes of distilled water using a portable power homogenizer (ASONE Corp., Osaka, Japan). The homogenates were processed according the kit’s instructions. The hydroxyproline signal (absorbance at 560 nm) was measured using a microplate reader (Spark^®^; Tecan Group, Ltd., Männedorf, Switzerland or SpectraMax; Molecular Devices, LLC., San Jose, CA).

### Histopathological analysis

Serial tissue sections were cut from paraffin-embedded lung specimens, and the first section (2-μm thick) was stained with H&E for histological examination and the remaining sections were used for immunohistochemcal analysis. The histopathological findings in this study for lung and mediastinal lymph node were determined after multifaceted discussions between certified pathologists from the Japanese Society of Toxicology and Pathology and certified medical pathologists from the Japanese Society of Pathology, based on the International Harmonization of Nomenclature and Diagnostic Criteria for Lesions in Rats and Mice (INHAND)[40]. Pathological diagnosis was performed blindly by three pathologists and summarized as a result of the discussion.

### Alcian blue staining

It is known that the alcian blue pH 1.0 method primarily detects sulfate groups, and the alcian blue pH 2.5 method preferentially detects carboxyl groups. Using both methods, CWAAP was stained blue by both staining methods. This study used the alcian blue pH 1.0 method, which did not stain the mucus of bronchial and alveolar epithelium (Fig. S4). After deparaffinization and rinsing, the slides were incubated in 0.1M HCl solution for 3 minutes. Then, they were incubated in alcian blue staining solution (alcian blue 8GX, C.I.74240, Merck-Millipore, Burlington, MA) for 10 minutes at room temperature. The slides were then lightly passed through a 0.1N HCl solution and washed with flowing water. Finally, after contrast staining with Kernechtrot (NUCLEAR FAST RED, C.I.60760, Merck-Millipore) for 5 minutes, the slides were processed for light microscopy.

### Masson’s Trichrome staining

The slides were deparaffinized, washed with water, and then reacted with a mixture of 10% potassium dichromate and 10% trichloroacetic acid for 60 minutes at room temperature. The specimens were then washed with water and stained with Weigelt’s iron hematoxylin solution (C.I.75290, Merck-Millipore) for 10 minutes at room temperature. The slides were then successively stained with 0.8% orange G solution (C.I.16230, Merck-Millipore) for 10 minutes at room temperature, Ponceau (C.I.14700, FUJIFILM-Wako Pure Chemical Corp., Osaka, Japan) acid fuchsin (C.I.42685, Merck-Millipore) azofloxine (C.I.18050, Chroma Germany GmbH, Augsburg, Germany) mixture for 40 minutes at room temperature, 2.5% phosphotungstic acid for 10 minutes at room temperature, and blue aniline solution (C.I.42755, Chroma Germany GmbH) for 10 minutes at room temperature. Between each staining the slides were washed lightly with 1% acetic acid water. The slides were then processed for light microscopy.

### PAS-Diastase staining

This method is a variant of PAS staining in which glycogen is degraded by pre-treatment with diastase to eliminate crossover against glycogen and improve specificity [41]. Slides were deparaffinized, rinsed with water, and digested with salivary amylase (in place of 1% diastase) at 37°C for 60 minutes. The slides were then washed in water, reacted with 0.5% periodate for 10 minutes at room temperature, washed with distilled water, and reacted with Schiff reagent (Cold Schiff’s reagent, #40932, Muto Pure Chemicals Co. Ltd., Tokyo, Japan) for 30 minutes at room temperature. The reaction was stopped by washing three times with sulfurous acid water. The slides were then washed with water and stained with Mayer’s Hematoxylin solution for 2 minutes at room temperature. After rinsing, the staining was checked under a microscope, and the slides were then processed for light microscopy.

### Immunohistological multiple staining analyses

Details of the multiple staining method have been described previously [42]. Briefly, lung tissue sections were deparaffinized with xylene, hydrated through a graded ethanol series, and incubated with 0.3% hydrogen peroxide for 10 min to block endogenous peroxidase activity. Slides were then incubated with 10% normal serum at room temperature (RT) for 10 min to block background staining, and then incubated for 2 hr at RT with the first primary antibody. After washing with PBS, the slides were incubated with histofine simple stain ratMAX-PO(MULTI) (414191, Nichirei, Tokyo, Japan) for 30 min at RT. After washing with PBS, slides were incubated with DAB EqV Peroxidase Substrate Kit, ImmPACT (SK-4103, Vector laboratories) for 2-5 min at RT. Importantly, after washing with dH_2_O after color detection, the sections were treated with citrate buffer at 98°C for 30 min before incubation with the next primary antibody to denature the antibodies already bound to the section. This procedure was repeated for the second and then the third primary antibodies. HighDef red IHC chromogen (ADI-950-142, Enzo Life Sciences, Inc., Farmingdale, NY) was used for the second coloration and Histogreen chromogen (AYS-E109, Cosmo Bio, Tokyo, Japan) for the third coloration. Coloration was followed by hematoxylin staining for 30-45 seconds. The slides were then processed for light microscopy. For immunofluorescence staining, all primary and secondary antibodies used were made into a cocktail for each staining step and used simultaneously. After the fluorescence-labeled secondary antibodies reaction, the sections were shielded with DAPI-containing encapsulant and used for imaging. The sections were observed under an optical microscope ECLIPSE Ni (Nikon Corp., Tokyo, Japan) or BZ-X810 (Keyence, Osaka, Japan). For measurement of phospho-SMAD3 and Tm4sf1 positive indices, the sham air group (n=5) and the 40 mg/m^3^ group immediately after exposure (n=7), and the sham air group (n=5) and the 40 mg/m^3^ group after an 18-week recovery period (n=8) were used for analysis. For the 40 mg/m^3^ exposure group, positive indexes were counted separately for multifocal lesions and normal surrounding tissue. In all animals, at least five fields of view were measured using a 40x objective lens. More than 500 cells per individual were measured for phospho-Smad3 and 300 cells per individual were measured for Tm4sf1, and the mean value per individual was used for statistical analysis.

### Statistical analysis

Except in the case of incidence and integrity of histopathological lesions, the data comparisons among multiple groups were performed by one-way analysis of variance with a post-hoc test (Dunnett’s or Tukey’s multiple comparison test), using GraphPad Prism 5 (GraphPad Software, San Diego, CA). The incidences and integrity of lesions were analyzed by the chi-square test using GraphPad Prism 5 (GraphPad Software, San Diego, CA). All statistical significance was set at *p* < 0.05.

### Availability of data and materials

The datasets used and analyzed during the current study are available from the corresponding authors on reasonable request.

## Abbreviations

ABCA3: ATP binding cassette subfamily A member 3
AEC1: Alveolar epithelial type 1 cell
AEC2: Alveolar epithelial type 2 cell
AEP: Alveolar epithelial progenitor
ARDS: acute respiratory distress syndrome
BAHyp: Bronchiolo-alveolar hyperplasia
BALF: Bronchoalveolar lavage fluid
CWAAP: Cross-linked water-soluble acrylic acid polymer
DAPI: 4’,6-diamino-2-phenylindole
EMT: Epithelial-Mesenchymal Transition
HE: Hematoxylin and eosin
LDH: Lactate dehydrogenase
LPCAT1: Lysophosphatidylcholine acyltransferase 1
MHLW: Ministry of Health, Labour and Welfare
MMAD: Mass median aerodynamic diameter
NBF: Neutral buffered formalin
PAS: Periodic acid Schiff
SAP: Super absorbent polymer
SEM: Scanning electron microscope
σg: geometric standard deviation
Smad3: Mothers against decapentaplegic homolog 3
Sox9: SRY-Box transcription factor 9
SP-D: Surfactant protein D
SUR: lesion-surrounding tissue
TGF: Transforming growth factor
Tm4sf1: Transmembrane 4 L six family member 1

## Acknowledgments

We would like to express our gratitude to the Collaborative Research External Evaluation Committee of the Japan Organization of Occupational Health and Safety and Hisao Yamaguchi, Manager, Masako Kiguchi, Director, Hirohide Ohnishi, Director and Dr. Shigeki Koda, Acting Director of the National Institute for Occupational Safety and Health, for their valuable advice in the planning and implementation of this study. In addition, we wish to thank Dr. David B. Alexander of Nanotoxicology project, Nagoya City University Graduate School of Medicine for his insightful comments and English editing. Moreover, we would like to express our sincere gratitude to the members of Matsuzawa Kosan and Total Service for their strong support in the breeding and dissection of the animals. Finally, we would like to express our heartfelt gratitude to all the Japan Bioassay Research Center staff.

## Author information

### Contributions

S.Y. and T.T. performed the experiments and analyzed the data. Y.G., K.M., S.H., Y.F., Y.K., K.M. and M.S. assisted with animal experiments. K.T., H.S., Y.U., and S.Y. performed histopathological diagnoses. S.Y., Y.U., Y.K., and K.O. conducted a pathological conference. M.S. and H.K. performed BALF sampling and dissection. Y.K. and K.O. and T.K. analyzed and interpreted the data. T.T. and S.Y. conceived, designed, and directed the study and interpreted the data. T.T., S.Y., and Y.U. drafted and revised the manuscript. All authors approved the manuscript as submitted.

### Ethics declarations

All animals were treated humanely and all procedures were performed in compliance with the Animal Experiment Committee of the Japan Bioassay Research Center.

## Supplementary Information

**Additional file 1: Figure S1.**
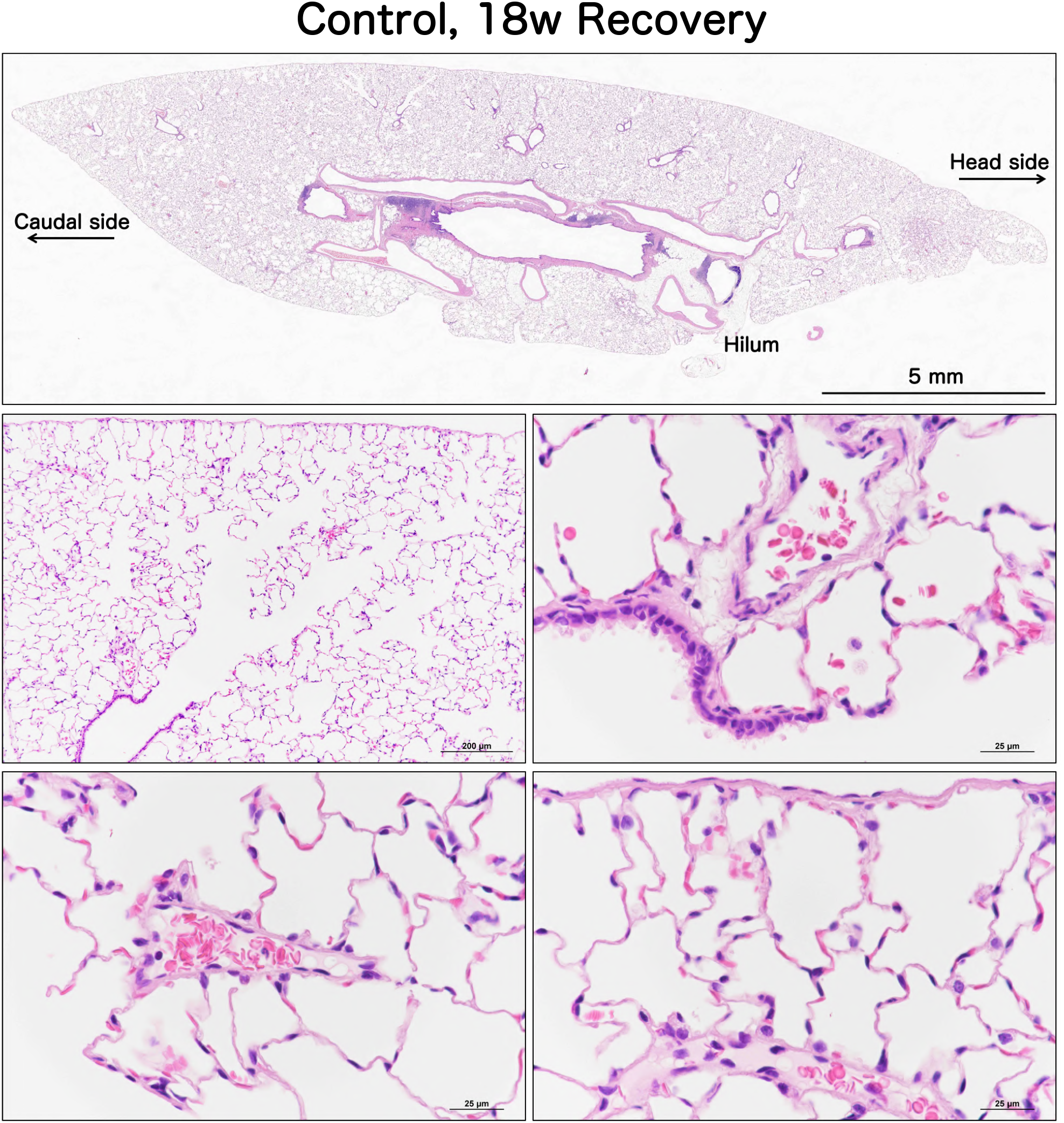
Representative microscopic photographs of a normal rat lung (sham air) after the 18 week recovery period.

**Additional file 2: Figure S2.**
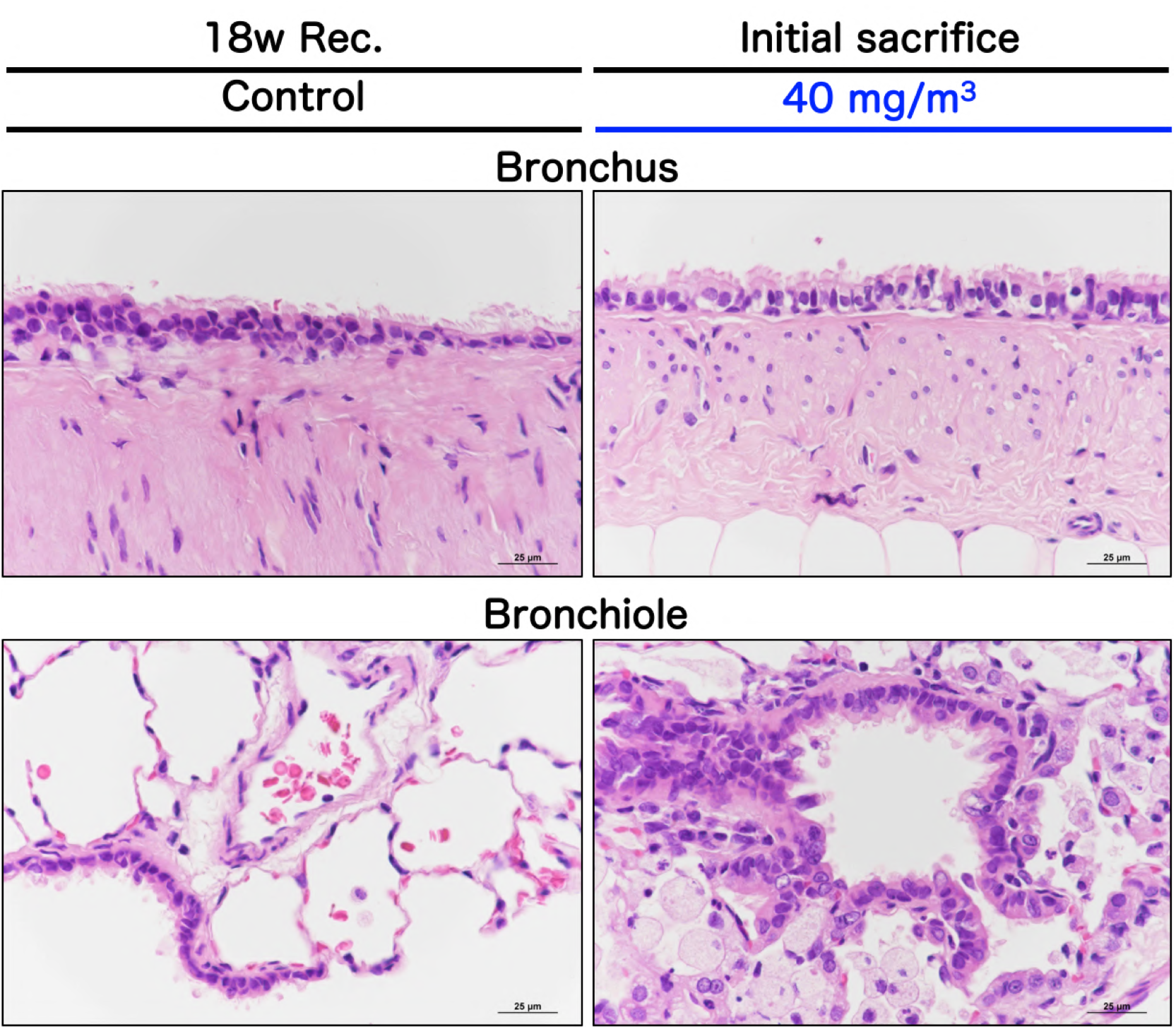
Representative histopathological photographs of the bronchus and bronchiole in the rat lung after repeated inhalation exposure to CWAAP-A (40 mg/m^3^).

**Additional file 3: Figure S3.**
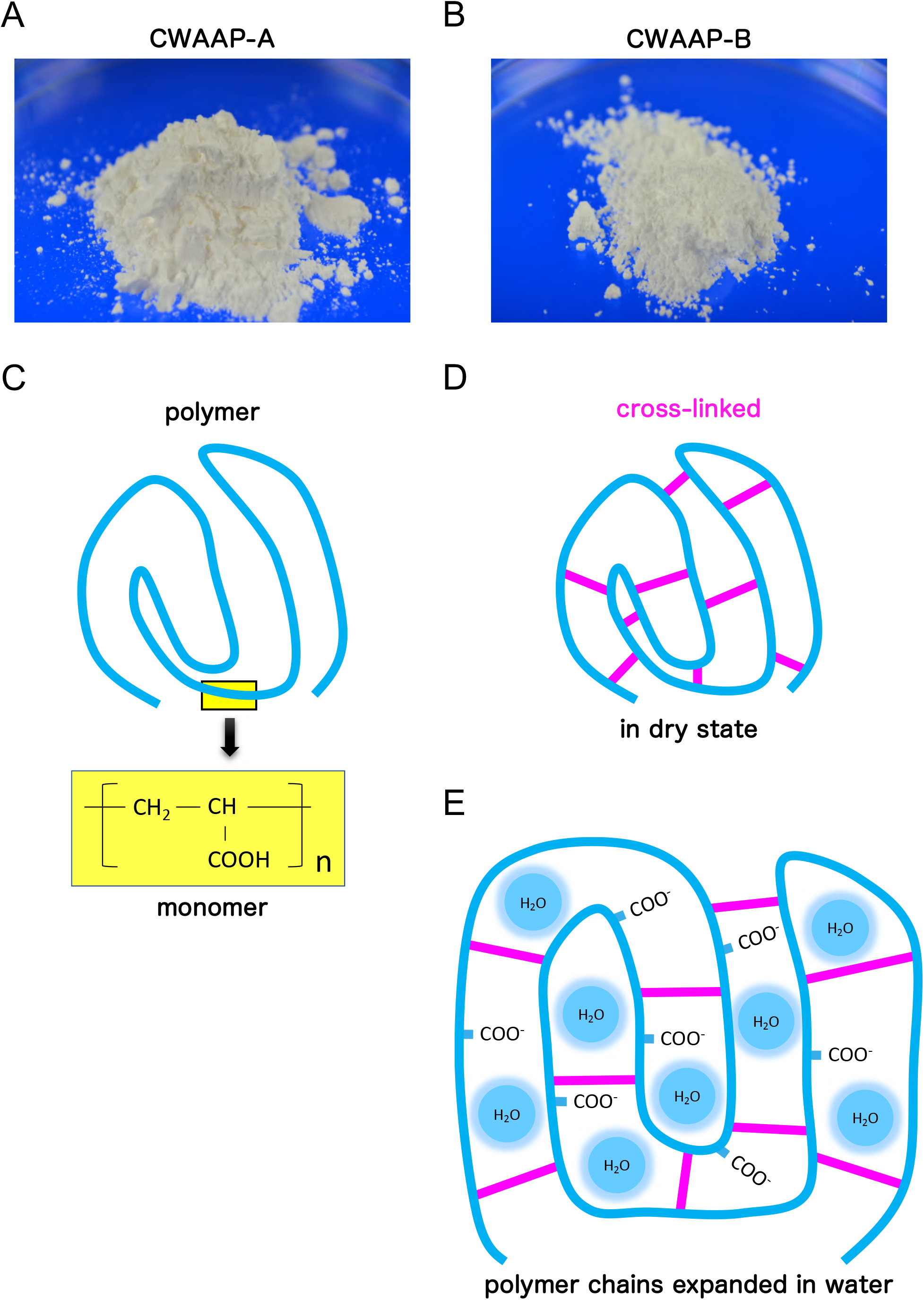
Representative images of CWAAP-A (A), CWAAP-B (B) and chemical structural formula of CWAAP (C-E). Acrylic acid polymer is a polymerized product of acrylic acid with carboxyl groups (C) and is anionic because of a large amount of carboxyl groups in the molecule. Cross-linked acrylic acid polymers (CWAAPs) have the characteristics of absorbing and retaining a large amount of water (E), as the polymer chain expands when it contains moisture compared to its dry state (D).

**Additional file 4: Figure S4.**
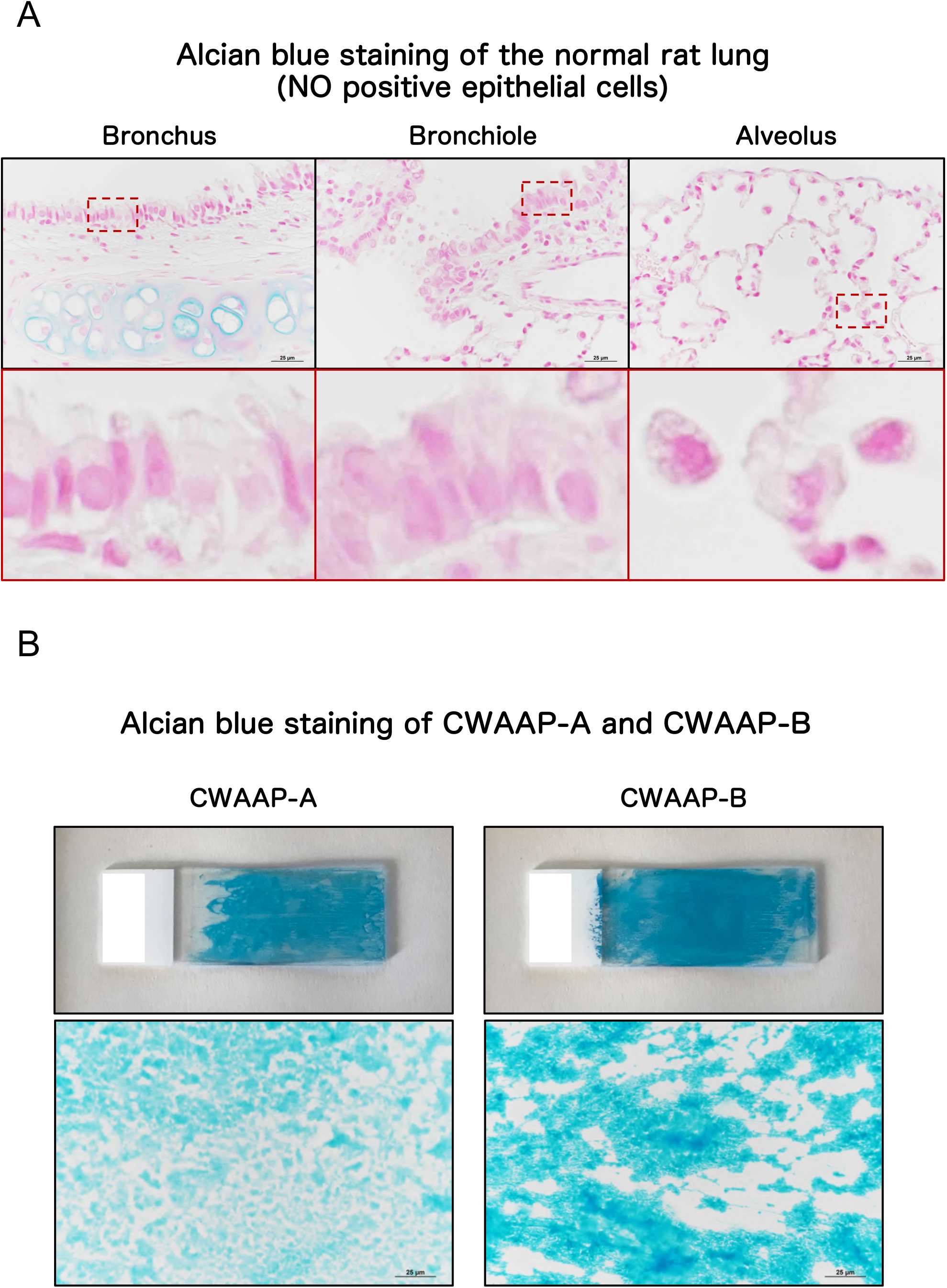
Representative images of the alcian blue staining of the normal rat lung are shown in A. In B, CWAAP-A and CWAAP-B were placed on the slides and directly stained using Alcian blue.

**Additional file 5: Figure S5.**
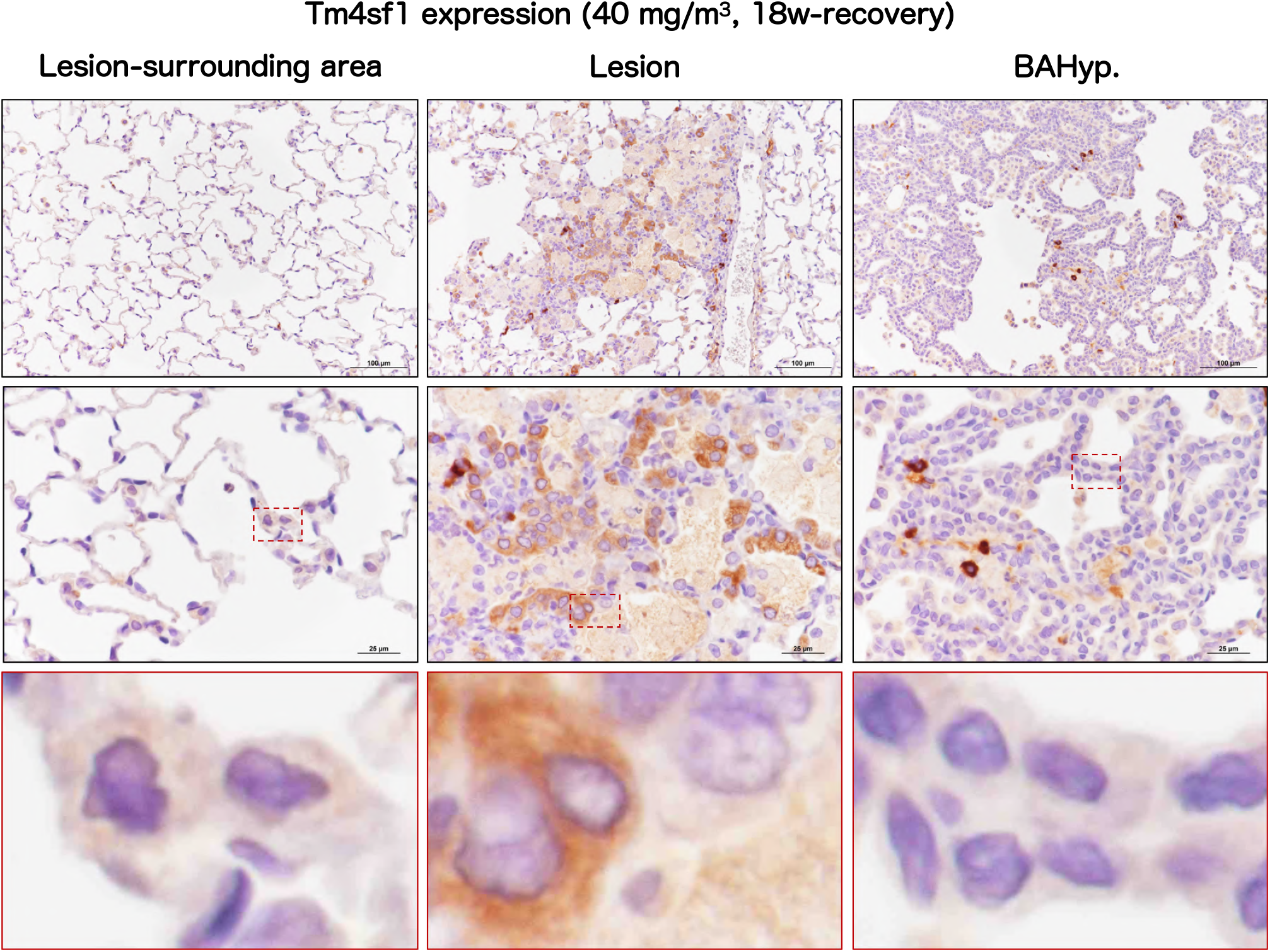
Representative images of Tm4sf1 expression in the rat lung after repeated inhalation exposure to 40 mg/m^3^ CWAAP-A.

**Additional file 6: Figure S6.**
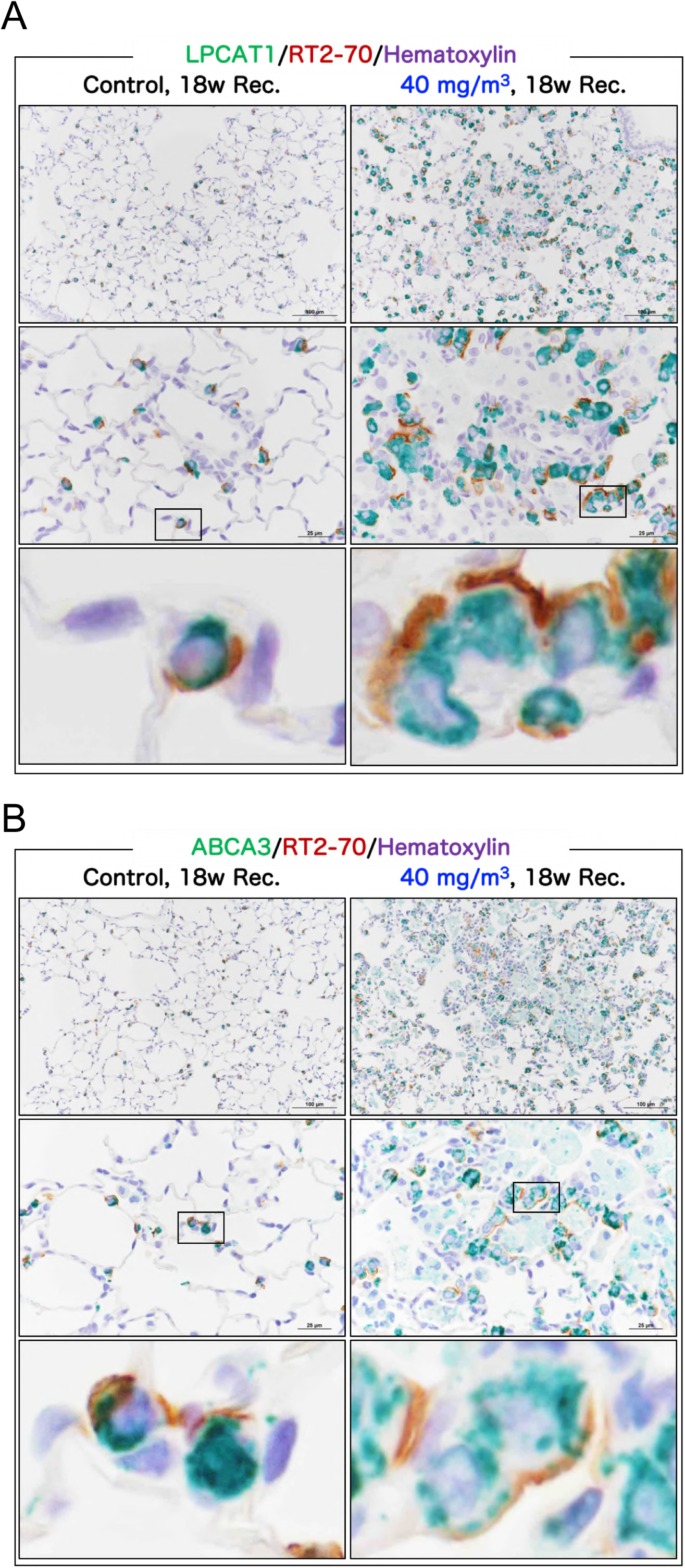
Representative images of the AEC2 membranous marker RT2-70 co-staining with AEC2 cytoplasmic markers LPCAT1 (A) and ABCA3 (B) in the rat lung.

**Additional file 7: Figure S7.**
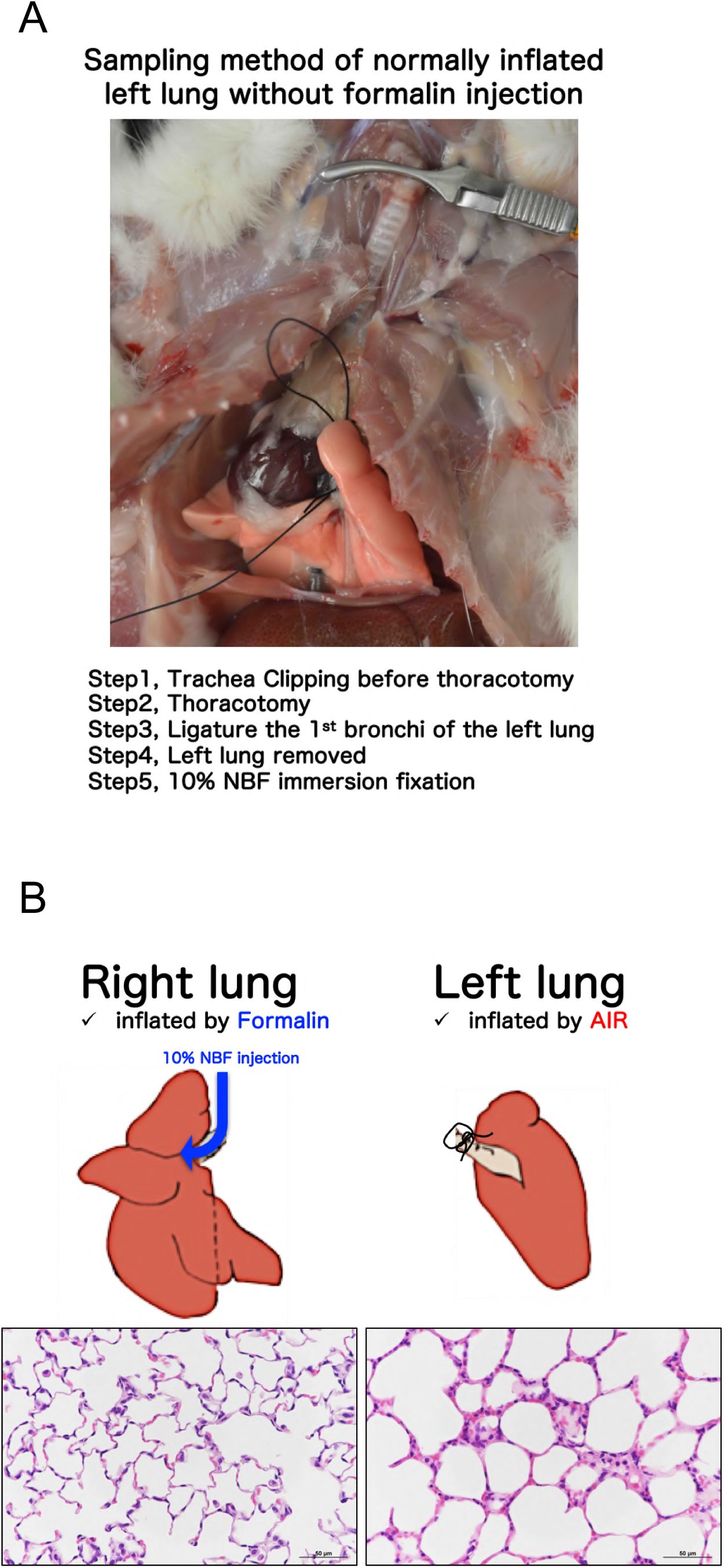
Sampling procedure of the air inflated left lung without formalin injection (A) and comparative images of the right lung inflated by formalin and the left lung inflated by air (B).

**Additional file 8: Figure S8.**
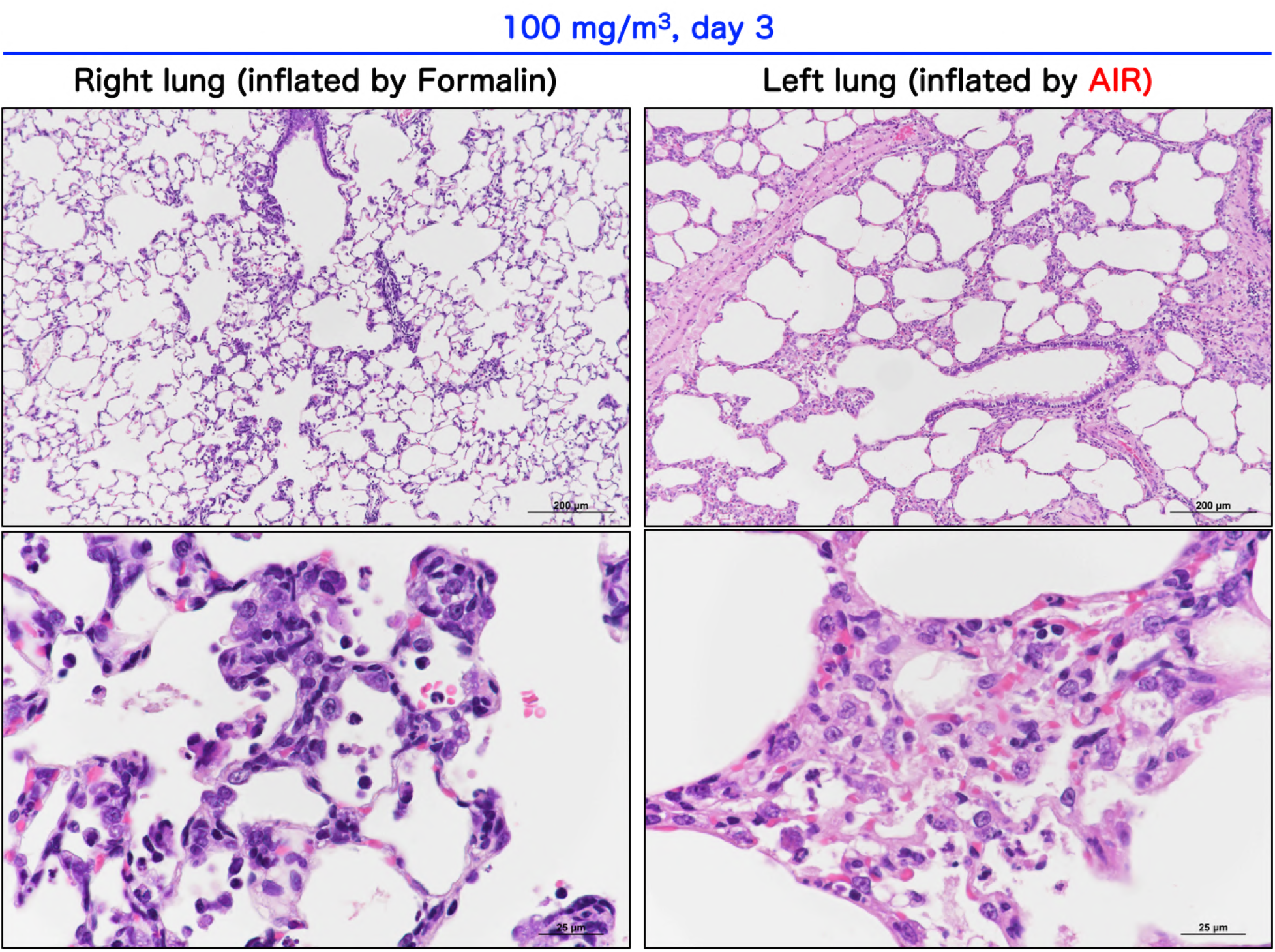
Representative histopathological images of the right lung inflated by formalin and the left lung inflated by air of a rat 3 days after exposure to 100 mg/m^3^: the right lung and the left lung are from the same animal.

**Additional file 9: Figure S9.**
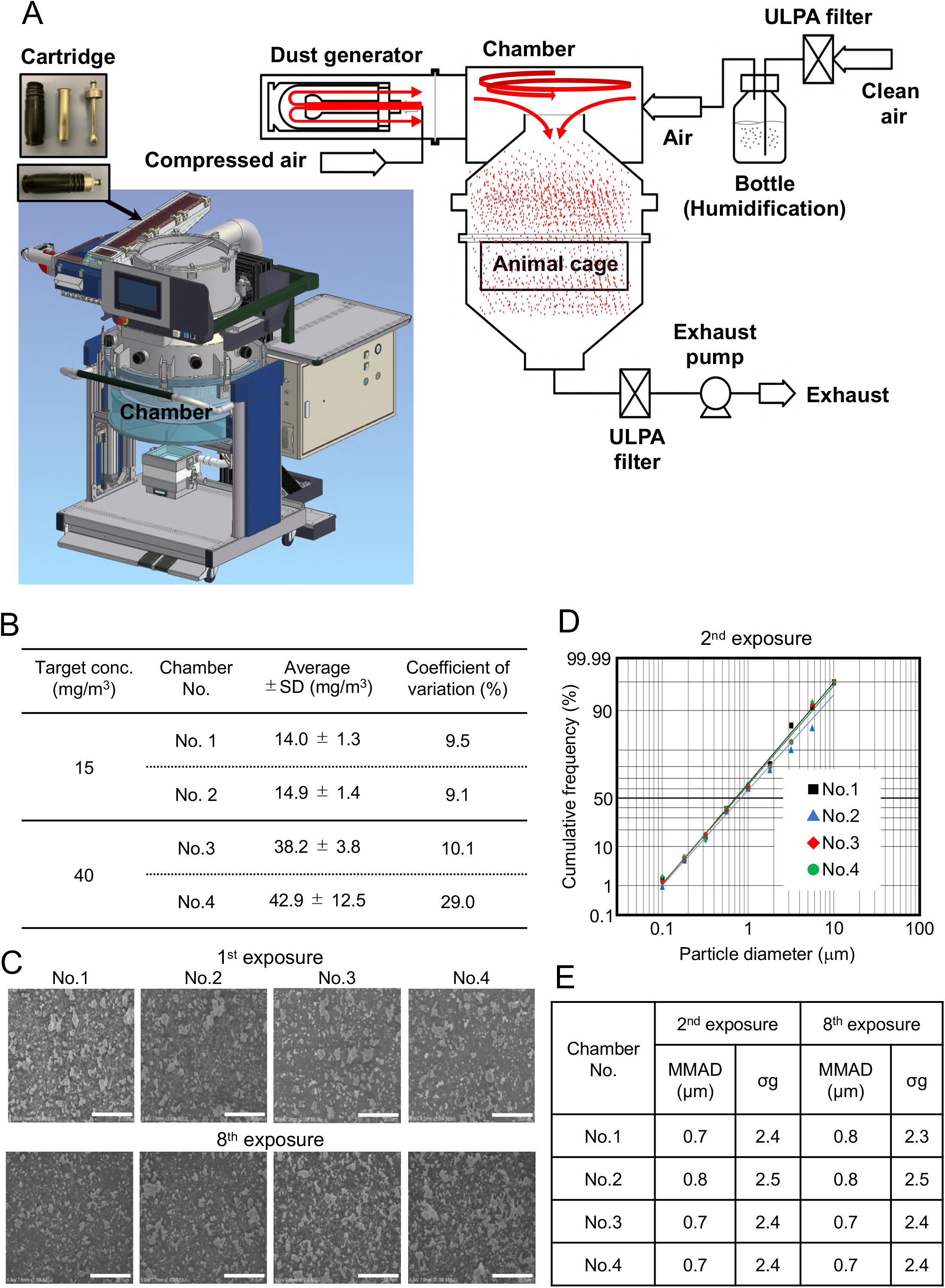
The direct-injection whole body inhalation system (A). Exposure concentrations of CWAAP-A in each chamber (B). Representative scanning electron microscope (SEM) images of the CWAAP particles in the chambers (C). Cumulative frequency distribution graphs with logarithmic probability (D). The mass median aerodynamic diameter (MMAD) and geometric standard deviation (σg) in the chambers measured during the second and eighth exposures (E). Scale bar: 20 μ

**Additional file 10: Figure S10.**
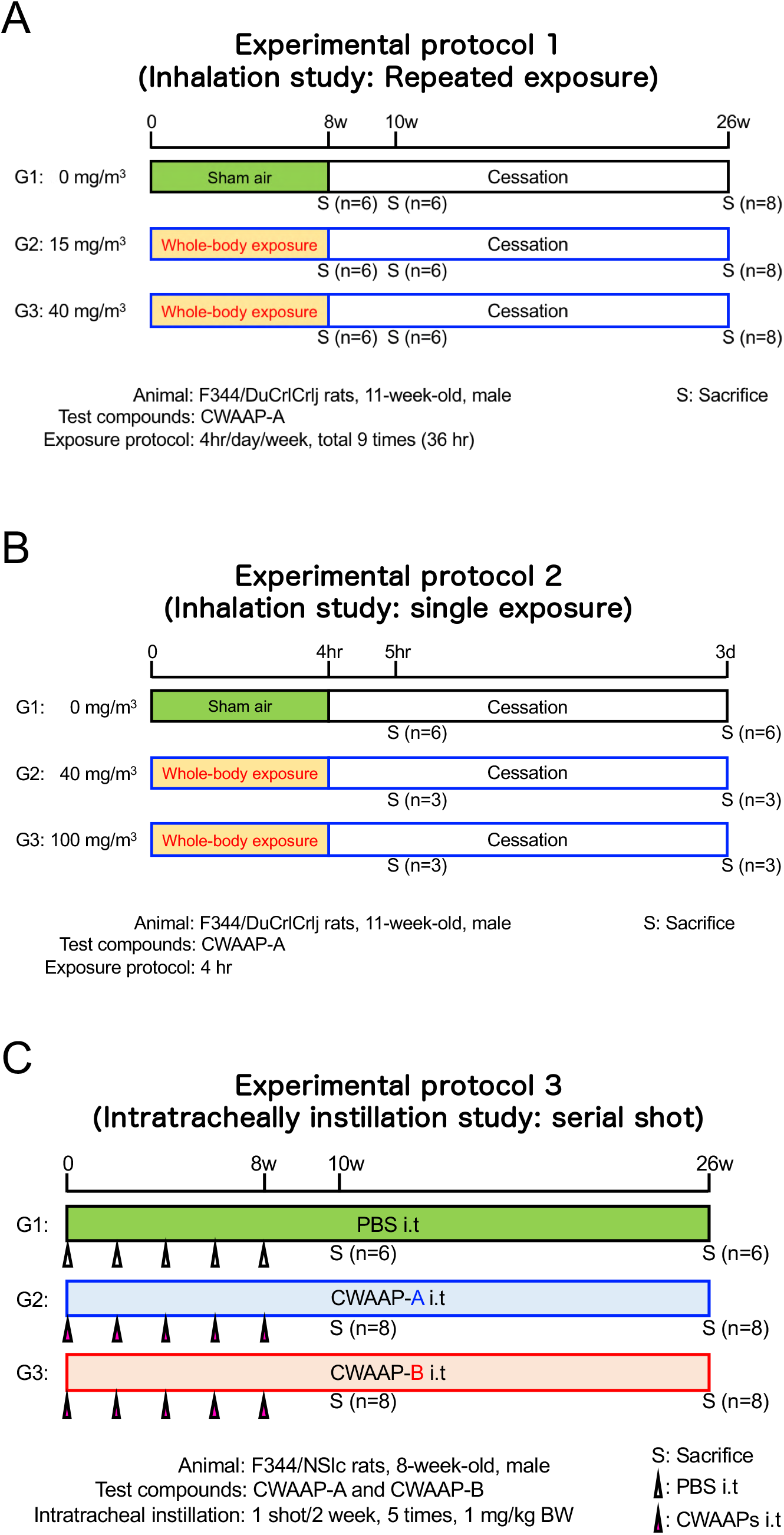
The three animal experimental protocols used in this study. The repeated inhalation exposure study (A), the single inhalation study (B), and the repeated intratracheal instillation study (C).

**Additional file 11: Figure S11.**
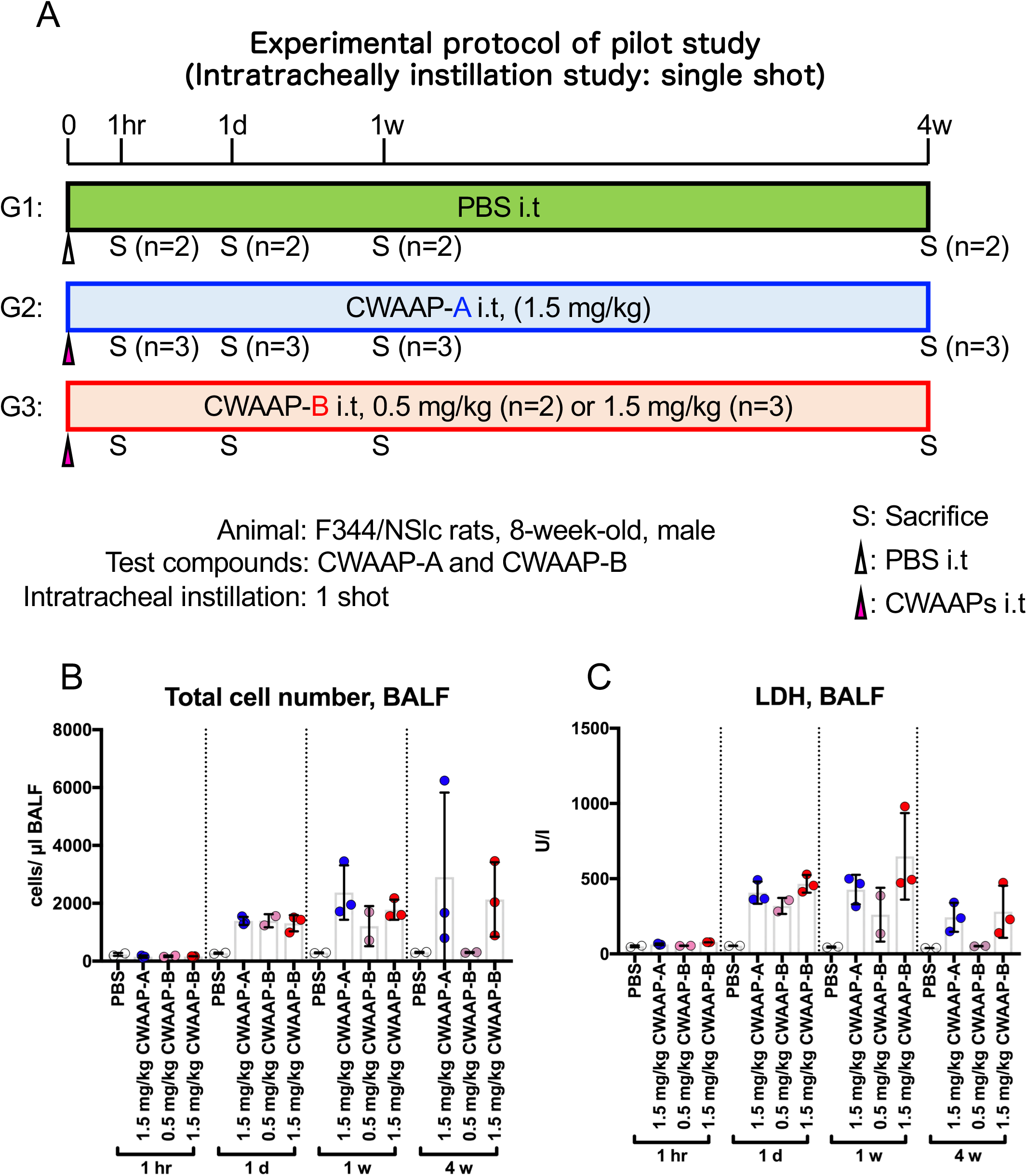
Experimental protocol of a pilot study using a single intratracheal instillation of 0.5 or 1.5 mg/kg CWAAP-A and CWAAP-B (A). Total cell number (B) and LDH activity (C) in the BALF of CWAAPs-treated rats and their respective controls (PBS).

**Additional file 12: Table S1.**
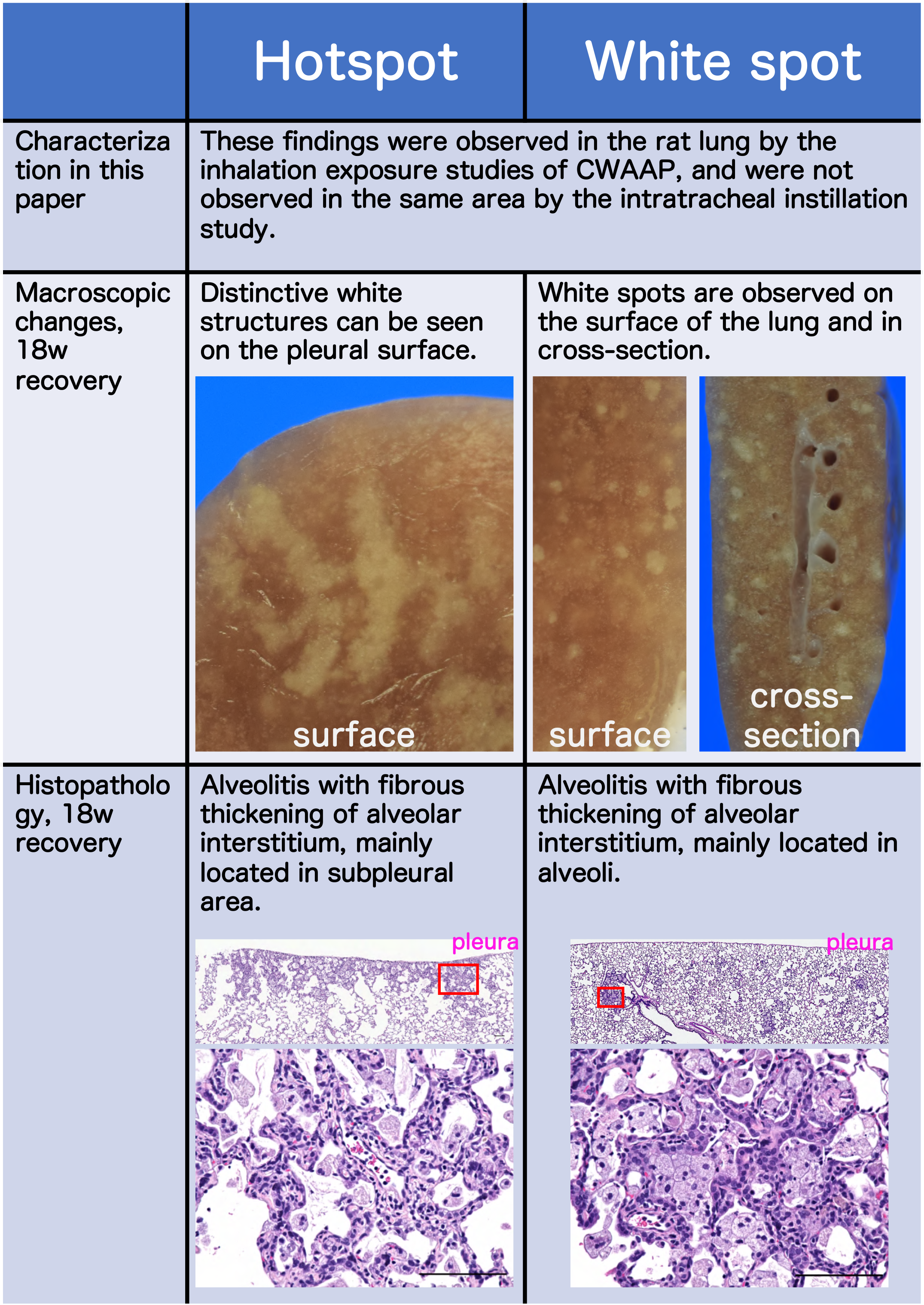
Summary of differences between hotspot and white spot.

**Additional file 13: Table S2.**
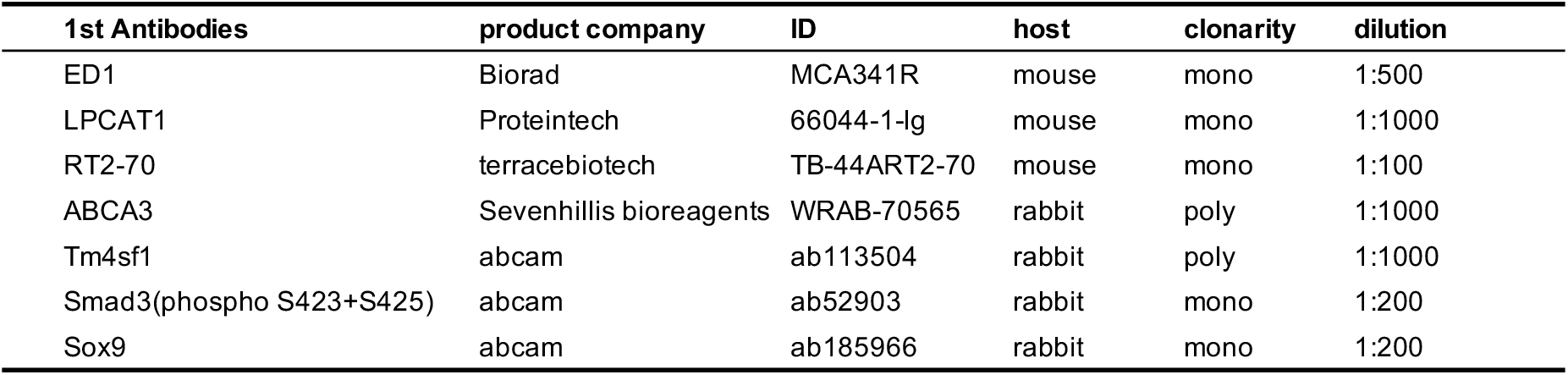
List of primary antibodies used in this study.

